# Molecular basis for differential activation of p101 and p84 complexes of PI3Kγ by Ras and GPCRs

**DOI:** 10.1101/2022.07.29.502076

**Authors:** Manoj K Rathinaswamy, Meredith L Jenkins, Xuxiao Zhang, Jordan TB Stariha, Harish Ranga-Prasad, Udit Dalwadi, Kaelin D. Fleming, Calvin K Yip, Roger L Williams, John E Burke

**Affiliations:** Department of Biochemistry and Microbiology, University of Victoria, Victoria, British Columbia, V8W 2Y2, Canada; MRC Laboratory of Molecular Biology, Cambridge, United Kingdom; Department of Biochemistry and Molecular Biology, The University of British Columbia, Vancouver, British Columbia V6T 1Z3, Canada

**Keywords:** PI3K, PIK3CG, PI3Kγ, p110γ, p101, p84, PIK3R5, PIK3R6, phosphoinositide 3-kinase, HDX-MS, Gβγ, GPCR

## Abstract

Class IB phosphoinositide 3-kinase (PI3Kγ) is activated in immune cells by diverse stimuli and can form two distinct complexes, with the p110γ catalytic subunit binding to either p101 or p84 regulatory subunits. These two complexes are differentially activated by G-protein coupled receptors (GPCRs) and Ras, but the molecular details of this activation are still unclear. Using a combination of X-ray crystallography, HDX-MS, EM, molecular modeling, and biochemical assays we reveal molecular differences between the two p110γ-p84 and p110γ-p101 complexes that explain their differential activation. The structure of p110γ-p84 shows a similar assembly to p110γ-p101 at the p110γ interface, however the interface in p110γ-p84 is dynamic and is evolutionarily conserved to be less stable compared to p110γ-p101. The p110γ-p84 complex is only weakly recruited to membranes by Gβγ subunits alone and requires recruitment by Ras to allow for Gβγ activation through an interaction with the p110γ helical domain. The interfaces of the p101 GBD with Gβγ, and the p110γ helical domain with Gβγ were determined using computational alphafold2 modeling and HDX-MS. There are distinct differences in the C-terminal domain of p84 and p101, which allows p101 to bind Gβγ subunits, while p84 does not. The two Gβγ interfaces in p110γ and p101 are distinct, revealing how unique mutants of Gβγ cause differential disruption of PI3Kγ complex activation. Overall, our work provides key insight into the molecular basis for how different PI3Kγ complexes are activated.

## Introduction

The class IB phosphoinositide 3-kinase PI3Kγ is a lipid kinase that generates the lipid signalling molecule phosphatidylinositol 3,4,5 trisphosphate (PIP_3_) downstream of diverse cell surface receptors (*1*). PI3Kγ can form two distinct complexes composed of a single catalytic subunit (p110γ, encoded by *PIK3CG*) binding to one of two regulatory subunits (p101 and p84, encoded by *PIK3R5* and *PIK3R6,* respectively) (*2–4*). PI3Kγ is highly expressed in immune cells, and is a master regulator of the adaptive and innate immune systems (*1*), with key roles in chemotaxis (*5*), reactive oxide production (*6*), and cytokine production (*7*). It also plays important roles in endothelial cells, neurons, cardiomyocytes, and pulmonary cells (*8*). Studies on catalytically dead PI3Kγ or using selective ATP-competitive inhibitors have defined important roles for it in the inflammatory response, and it shows promise as a therapeutic target for inflammatory disease including lupus (*9*), arthritis (*10*), atherosclerosis (*11*), asthma (*12*), and obesity related changes in metabolism (*13, 14*). Overexpression of *PIK3CG* is observed in cancer (*15, 16*), and targeting PI3Kγ as an immunomodulator of the tumor-microenvironment has shown promise as an anti-cancer therapeutic (*17, 18*), with PI3Kγ selective inhibitors in phase II clinical trials for triple negative breast cancer, renal cell-carcinoma, and urothelial carcinoma (*19*). However, the discovery of primary immunodeficiency patients harboring loss of function mutations in PI3Kγ (*20, 21*) highlights potential challenges in therapeutic inhibition.

The two complexes of PI3Kγ (p110γ-p101 and p110γ-p84) play unique roles in cellular processes, with these putatively mediated by their differential ability to be activated by diverse stimuli, including G-protein coupled receptors (GPCRs) (*22*), the IgE/Antigen receptor (*6*), receptor tyrosine kinases (*23*), and Toll-like receptors (TLRs) (*24*). Experiments examining immune cells with selective knockout of the p101 or p84 regulatory subunits show that p101 is required for PI3Kγ’s role in chemotaxis, while the p84 subunit is required for reactive oxide generation (*25–27*), with knockout of both regulatory subunits leading to complete loss of PI3Kγ activity (*26*). Biochemical reconstitution studies have defined two major signaling proteins that mediate PI3Kγ activation downstream of cell surface receptors, lipidated Gβγ subunits released by activated GPCRs, and GTP loaded lipidated Ras. The presence of p101 and p84 regulatory subunits dramatically alter the activation by each of these stimuli, with *in vitro* the p110γ-p101 complex is activated ∼100 fold by Gβγ, while p110γ-p84 is activated ∼5 fold (*28–32*). In cells the p110γ-p84 complex is poorly recruited to cell membranes by Gβγ subunits, with it requiring Ras for membrane localization (*32*). The p101 subunit forms an obligate heterodimer with p110γ, while p84 forms a weaker transient interaction with p110γ (*31*), but the molecular basis for this is currently not understood.

Extensive biophysical experiments on the free p110γ catalytic subunit and the p110γ-p101 complex have revealed insight into the architecture and regulation of p110γ (*28, 30, 33–35*). The p110γ catalytic subunit is composed of an adaptor binding subunit (ABD), a Ras binding domain (RBD) that mediates activation downstream of Ras, a C2 domain, a helical domain, and a bilobal kinase domain. The cryo-EM structure of the p110γ-p101 complex revealed that p110γ binds to the p101 regulatory subunit through the C2 domain, and the RBD-C2 and C2-helical linkers (*30*). We previously mapped a putative Gβγ binding interface in the helical domain of p110γ (*28*), with an additional binding site in the C-terminal domain of p101 (*28, 30*). Mutations in Gβγ have differential effects on either p110γ or p110γ-p101 activation (*36*), but the full molecular details of how Gβγ binds to either p110γ or p101 is still unclear.

To decipher the molecular mechanism for why p101 and p84 subunits differentially regulate p110γ activation, we determined the structure of the p110γ-p84 complex using a combined X-ray crystallography, EM, and computational modeling approach. Hydrogen deuterium exchange mass spectrometry (HDX-MS) experiments revealed that the p110γ-p84 is dynamic relative to the p110γ-p101 complex. Membrane reconstitution experiments using HDX-MS to study membrane recruitment of p110γ-p84 mediated by lipidated Gβγ and Ras shows that p110γ-p84 requires Ras for membrane localization. The p110γ-p84 complex can only be potently activated by Gβγ when Ras is present, where the p110γ-p101 complex can be robustly activated by Gβγ subunits alone. Finally, computational modeling and HDX-MS were used to define the Gβγ binding interfaces with both the C-terminal domain of p101 and the helical domain of p110γ. Overall, this work provides unique insight into the molecular mechanisms mediating differential PI3Kγ activation by Ras and GPCR signalling.

## Results

### Structure of the p110γ-p84 complex

To understand differences in the regulation of p110γ-p84 versus p110γ-p101 required molecular details of the p110γ-p84 complex. We purified full length human p110γ in complex with either mouse p84 or porcine p101, with gel filtration profiles consistent with the formation of heterodimers. The domain architecture of p110γ, p84, p101 are shown in Fig 1A, and the full list of all proteins and protein complexes purified in this manuscript shown in Fig. S1. To determine the structure of the p110γ-p84 complex we utilised a combination of X-ray crystallography, electron microscopy (EM), and Alphafold2 computational modelling. Initial negative stain EM data revealed that purified p110γ-p84 was homogenous and formed a similar shaped complex to our recently determined p110γ-p101 Cryo-EM complex. However, even with extensive optimisation we could not generate high quality vitrified specimens for Cryo-EM, as the p110γ-p84 complex always dissociated into free p110γ and p84 particles. Extensive screening of precipitant conditions allowed us to obtain crystals of p110γ-p84 that diffracted to ∼8.5 Å, with initial attempts to phase this using molecular replacement with the p110γ-p101 cryo-EM structure being unsuccessful. To provide additional molecular details on this complex we utilised an Alphafold2 (*37*) model specifically trained for multimeric complexes (*38*). Extensive computational modelling of different sequences of p110γ and p84 resulted in a consensus solution for the interface of p110γ with p84. These models had low predicted alignment error (PAE) between the p110γ and p84 subunits, which is a measure of the confidence of protein-protein interfaces (Fig S2). This model was then used as a search model for the low-resolution X-ray diffraction data, with only rigid body refinement resulting in a solution with high confidence (rwork=0.28, rfree=0.34, Table S1) despite the low resolution of the X-ray diffraction (Fig. 1C, S3). While the absolute positioning of side chains is challenging at this resolution, analysis of the 2fo-fc density revealed the orientation of the helices in p84 at the p110γ interface, validating the inter-subunit orientation (Fig. S3B), with this solution fitting well in the low resolution negative stain EM density (Fig. S3A).

**Figure 1.**
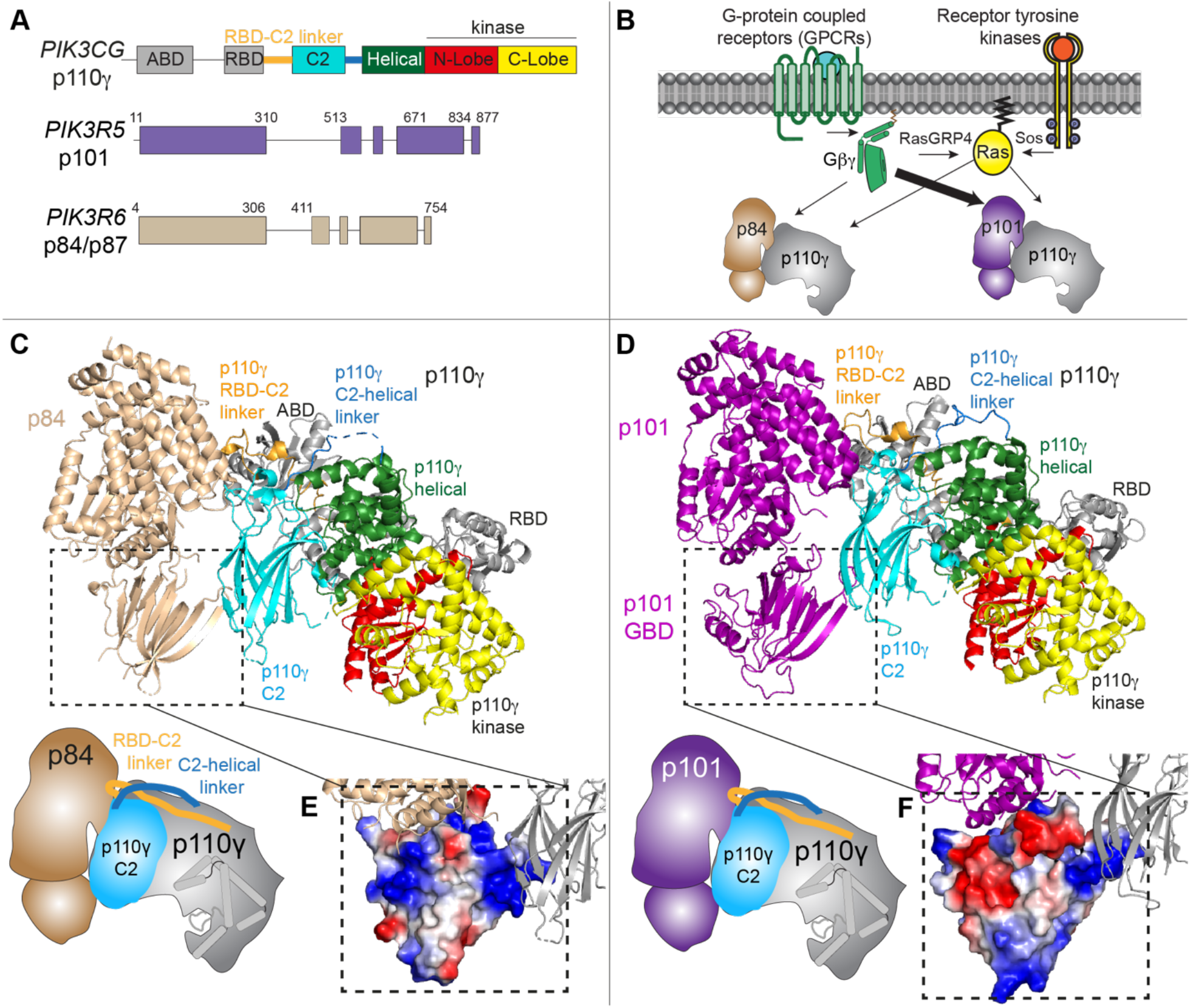
The structure of the p110γ-p84 complex and comparison with p110γ-p101. **A.** Cartoon schematic of the PI3Kγ catalytic (p110γ) and regulatory subunits (p101 and p84) with domain boundaries indicated. **B.** Cartoon of differences in activation between p110γ-p84 and p110γ-p101 complexes downstream of GPCRs and RTKs. **C.** Model of the p110γ-p84 complex based on X-ray crystallography, negative stain EM and alphafold modelling (Fig. S2+S3). Domains are indicated from panel A, with a cartoon schematic shown in the bottom left. **D.** Structure of the p110γ-p101 complex (PDB: 7MEZ) (*30*). Domains are indicated from panel A, with a cartoon schematic shown in the bottom left. **E+F.** Differences in the C-terminal domain of p84 (**E**) and p101 (**F**) are shown with this domain shown as an electrostatic surface.

The overall architecture of the p84 subunit is conserved compared to p101, with it containing a N-terminal helical domain, a central α/β barrel domain, and a C-terminal beta sandwich domain. The orientation of the regulatory subunit in the p110γ-p84 complex versus the p110γ-p101 complex (*30*) was strikingly similar (Fig. 1C+1D), with p84 binding to the C2 domain and the RBD-C2 and C2-helical linkers of p110γ. The primary interface for p110γ in p84 was located at the N-terminal helical region of p84, with additional interactions involving the GBD and C-terminus. One of the primary differences between p101 and p84 regulatory subunits is their differential ability to be recruited by lipidated Gβγ subunits. We have previously identified that this binding occurs at the C-terminal domain of p101, in a region we defined as the Gβγ binding domain (GBD) (*28, 30*). The Alphafold2 model of the p110γ-p84 structure allowed us to examine differences in this domain. The C-terminal domain of p84 contains the same β-sandwich fold, however, there are distinct differences compared to p101 at the face of this domain distal from p110γ. This can be clearly highlighted by visualizing the electrostatics of the GBD between p101 and p84, showing a strikingly different interfacial surface for Gβγ binding (Fig 1E+F). One of the primary differences that was immediately apparent was the presence of a helical extension in the C-terminal domain of p101 that is part of the Gβγ binding face (*28, 30*) which is not present in p84.

### Differences in the interface of p84 with p110γ compared to p101

Previous *in vitro* assays testing subunit exchange of p110γ-p101 and p110γ-p84 complexes suggest that the p101 complex forms a constitutive complex with p110γ, with the p84 complex forming a weaker dynamic interaction with p110γ (*31*). The structures of p110γ-p84 and p110γ-101 reveal that the regulatory subunits bind to the same interface with p110γ. The most extensive binding interface for both p84 and p101 with p110γ is comprised of a set of N-terminal helices, specifically the loops between α3-α4 and α5-α6 (Fig. 2A+B). In both p84 and p101 the interfacial residues found in these helices are strongly conserved across evolution (Fig. 2C), however, there are distinct differences between the two subunits. Within the p110γ-101 complex we had previously identified a set of cation-pi interactions between charged residues in p110γ and aromatic residues in p101 (*30*). This includes the interaction between R362 in p110γ with Y75 in p101 and W413 in p110γ, and the interaction between R359 in p110γ and W115 in p101. These cation-pi interactions are lost in p84, with the conserved Y75 in p101 being replaced by a conserved Arg or Lys residue in the corresponding position in p84, and W115 in p101 being replaced by L108 in the corresponding position in p84 (Fig. 2C).

**Figure 2.**
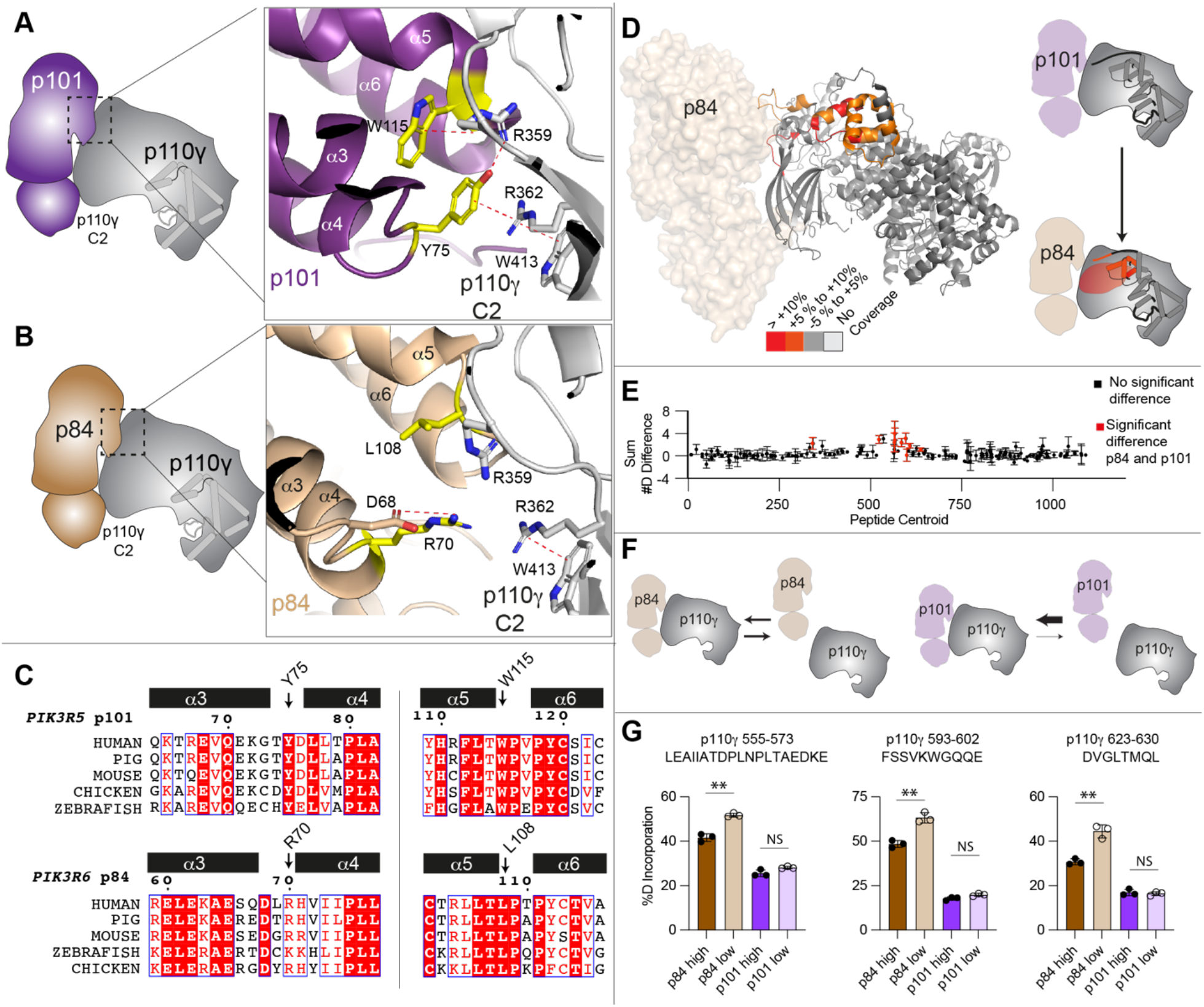
The p110γ interface is weaker in p84 versus p101. **A+B**. Cartoon schematic of the p110γ interface for p101 (A) and p84 (B), with a zoom in on the residues localised interface between both p84 and p101 with p110γ.Dotted lines indicate cation-pi or electrostatic interactions. **C.** Sequence alignment of both p101 and p84 residues in the α3 to α6 helices located at the p110γ interface. The residues annotated in panel are indicated on the alignment. **D.** HDX-MS differences in the p110γ subunit between the p110γ-p101 and p110γ-p84 complex. Significant differences in deuterium exchange (defined as greater than 5%, 0.4 Da, and a two-tailed t-test p<0.01 at any timepoint are mapped on to the structure of p110γ-p84 and cartoon of p110γ according to the legend. **E.** Sum of the number of deuteron difference between the p110γ-p101 and p110γ-p84 complexes over the entire deuterium exchange time course. Positive difference is indicative of enhanced exchange in p110γ-p84. Each point is representative of the centre residue of an individual peptide. Peptides that met the significance criteria described in C are coloured red. Error is shown as standard deviation (n=3). All HDX-MS data is provided in the source data. **F.** Cartoon schematic of the proposed equilibrium for dissociation of the two complexes. **G.** Selected deuterium exchange at 30 seconds for peptides in p110γ for p110γ-p101 and p110γ-p84 complexes at either high concentration (1500 nM) or low concentration (175 nM). Error is shown as standard deviation (n=3) with two tailed p-values as indicated: **<0.01; not significant (ns)>0.05.

To further define conformational differences between the two complexes we carried out experiments using HDX-MS, which measures dynamic differences in secondary structure in proteins. These experiments were carried out with human p110γ bound to porcine p101 and human p110γ bound to mouse p84. We compared the H/D exchange rates in p110γ between the two complexes at four different time points (3s, 30s, 300s and 3000s). The full details of HDX-MS data processing are in Table S2, with all raw HDX-MS data for all time points available in the source data. We observed statistically significantly decreased exchange (defined as differences at any time point >5%, >0.4 Da, and a p-value less than 0.01) in the p110γ-p101 complex versus the p110γ-p84 complex in the helical domain, C2 domain, the RBD-C2 linker and the C2-helical linker (Fig. 2D+E). These changes were all localised to either the interface with regulatory subunits, or the helical domain adjacent to the interface, which is consistent with p101 forming a more stable complex with p110γ compared to p84.

To further validate the dynamic nature of the p110γ-p84 complex compared to the p110γ-p101 complex we carried out HDX-MS experiments with varying concentrations of p110γ-p84 or p110γ-p101. In addition, for these experiments we purified p110γ bound to either the human p84 or p101 regulatory subunits, to validate that the changes observed were not due to minor differences in evolutionary conservation between mouse, pig, and human sequences. HDX-MS experiments were carried out at two timepoints (30 and 300 sec) with a final concentration of 1500 nM in high concentration experiments and 175 nM in low concentration experiments for both p110γ-p84 and p110γ-p101. Comparing p110γ-p84 with p110γ-p101 in the high concentration experiment showed similar differences to what we observed with the mouse p84 or pig p101 complexes, showing the difference between regulatory subunits is conserved for the human proteins (source data). For the p110γ-p101 complex there was no significant difference in exchange between the high and low concentration samples, signifying that the complex remains intact in both conditions (Fig. 2G). However, in the p110γ-p84 complex there was significant increases in exchange at p84 interfacial regions in the low concentration compared to high concentration (Fig. 2G). This is consistent with the p84 complex being dynamic compared to p110γ-p101 (Fig. 2F).

### Activation of the p110γ-p84 / p110γ-p101 complexes by lipidated Gβγ and Ras

To provide additional insight into functional differences between p110γ-p84 and p110γ-p101 we characterised their lipid kinase activities as well as the activity of the free p110γ catalytic subunit, using membrane reconstitution assays with lipidated Gβγ and lipidated GTPγS loaded G12V HRas (Fig. 3A). We characterized the lipid kinase activities using saturating concentrations of lipidated Gβγ and lipidated HRas on membranes roughly mimicking the composition of the plasma membrane (5% PIP_2_, 20% phosphatidylserine, 50% phosphatidylethanolamine, 10% cholesterol, 10% phosphatidylcholine, and 5% sphingomyelin). The presence of HRas alone led to roughly similar 3-fold activation for p110γ, p110γ-p84, and p110γ-p101 (Fig. 3B). The presence of Gβγ alone led to robust activation of p110γ-p101 (>100 fold activation), with weak activation of p110γ-p84 (∼3 fold), and no detectable activation of p110γ (Fig. 3B). The additional presence of HRas for p110γ-p101 with Gβγ caused an approximately similar 3-fold activation as was seen in the absence of Gβγ. However, for both free p110γ and p110γ-p84 there was a large synergistic activation when both HRas and Gβγ were present. This was consistent with previous observations of Gβγ and HRas activation on other membrane systems (*28–30, 32*). Because the p110γ-p84 complex is more reliant on activation by Ras, we wanted to ensure that there was no major affinity difference towards HRas for p110γ-p84 and p110γ-p101. We carried out activation assays with varying levels of HRas, both in the presence and absence of saturating lipidated Gβγ subunits. Both p110γ-p84 and p110γ-p101 in the presence and absence of Gβγ showed very similar EC50 values (Fig. 3C+D).

**Figure 3.**
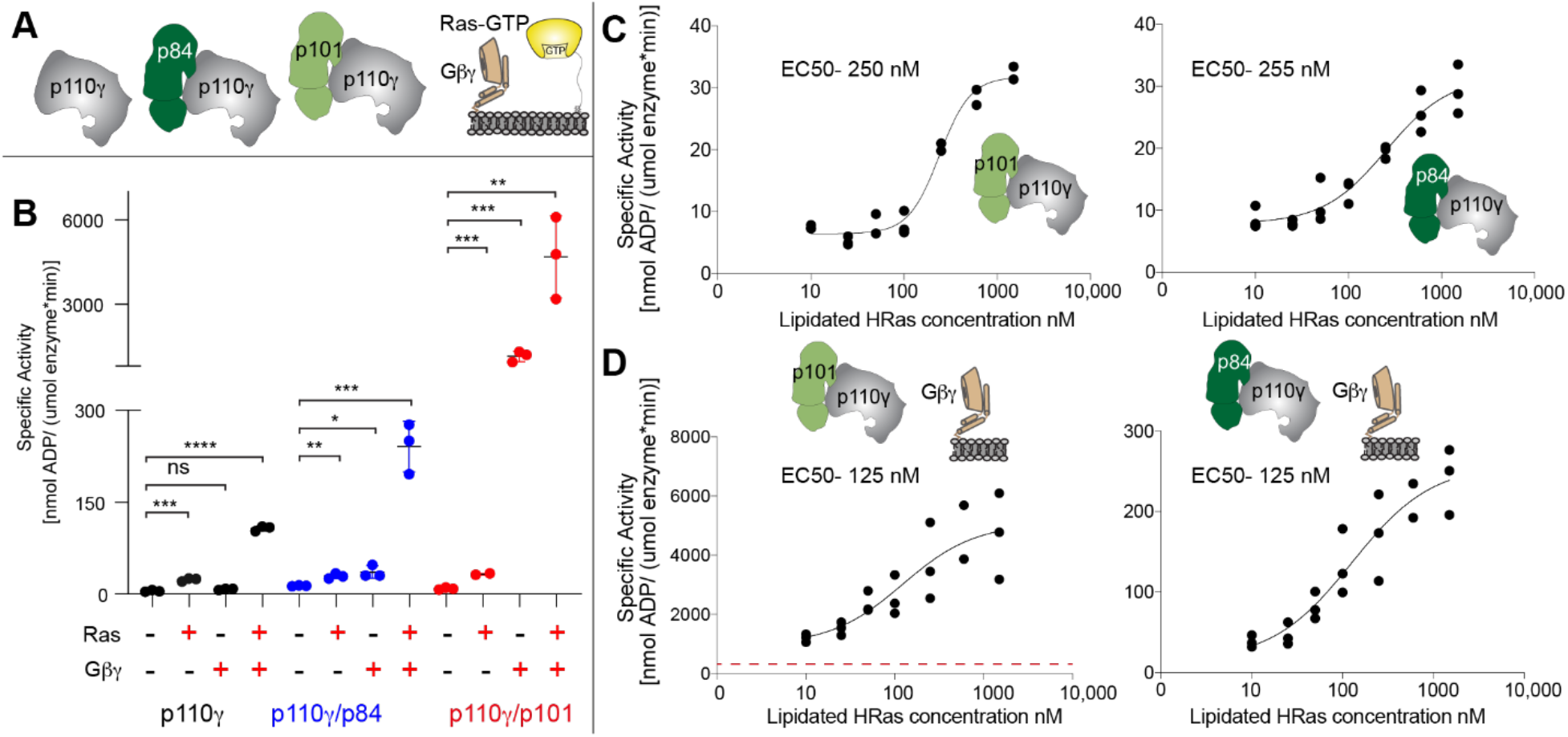
Activation of p110γ-p84 and p110γ-p101 by lipidated HRas and Gβγ. **A.** Cartoon schematic describing PI3Kγ variants tested and the lipidated activators, GTPγS loaded HRas and Gβγ. **B.** Lipid kinase activity assays of different p110γ complexes (concentration, 100 to 2000 nM) with and without lipidated Gβγ (1.5 μM) and lipidated HRas (1.5 μM) using 5% phosphatidylinositol 4,5-bisphosphate (PIP_2_) vesicles mimicking the plasma membrane (20% phosphatidylserine, 50% phosphatidylethanolamine, 10% cholesterol, 10% phosphatidylcholine, 5% sphingomyelin). Error bars represent standard deviation (n=3). Two tailed p-values represented by the symbols are as follows: ****<0.0001, ***<0.001; **<0.01; *<0.05; not significant (ns)>0.05 **C.** Lipid kinase activity assays of p110γ-p84 and p110γ-p101 with varying concentrations of lipidated HRas. **D.** Lipid kinase activity assays of p110γ-p84 and p110γ-p101 in the presence of lipidated Gβγ (1.5 μM) with varying concentrations of lipidated HRas. Experiments in panels C+D were performed using the same vesicles as in panel B. The dotted red line in the graph for the p110γ-p101 complex shows the peak activity for p110γ-p84 with both activators.

### HDX-MS analysis of Gβγ and HRas activation of p110γ-p84

To define the molecular mechanism underlying the difference between p110γ-p84 and p110γ-p101 activation by lipidated Gβγ and HRas we carried out HDX-MS experiments on membrane reconstituted complexes. HDX experiments were carried out for four time points (3s, 30s, 300s, 3000s) for five conditions: p110γ-p84 alone, p110γ-p84 with PM mimic vesicles, p110γ-p84 with HRas on PM mimic vesicles, p110γ-p84 with Gβγ on PM mimic vesicles, and p110γ-p84 with Gβγ and HRas on PM mimic vesicles (Fig. 4). The full details of HDX-MS data processing are in Table S2, with all raw HDX-MS data for all time points available in the source data.

**Figure 4.**
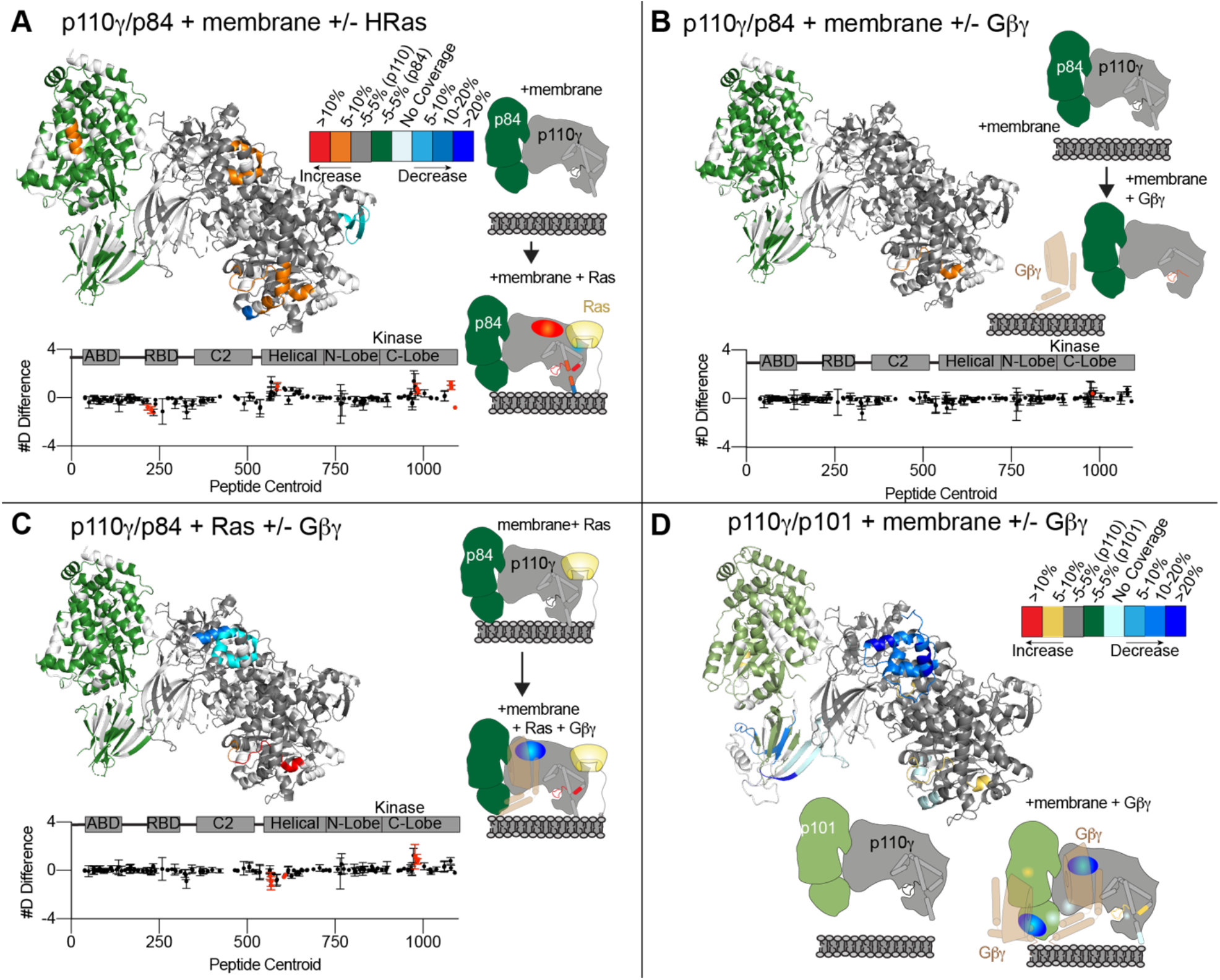
HDX-MS analysis of p110γ-p84 activation by membrane localised HRas and Gβγ, and comparison to p110γ-p101. **A-C.** Significant HDX-MS differences in the p110γ and p84 subunits between **(A)** plasma membrane mimic vesicles and plasma membrane mimic vesicles with 3 μM GTPγS loaded lipidated HRas **(B)** plasma membrane mimic vesicles and plasma membrane mimic vesicles with 3 μM Gβγ **(C)** plasma membrane mimic vesicles with 3 μM GTPγS loaded lipidated HRas and plasma membrane mimic vesicles with both HRas and Gβγ (3 μM) are mapped on the structure of p110γ-p84 according to the legend in panel A. A cartoon model is shown to the right with differences annotated. The sum of the number of deuteron difference is shown for p110γ, with red dots representing peptides showing statistically significant differences. **D.** Significant HDX-MS differences in the p110γ and p101 subunits between plasma membrane mimic vesicles and plasma membrane mimic vesicles with Gβγ mapped on the structure of p110γ-p101 (PDB:7MEZ) according to the legend. A cartoon model is shown to the right with differences annotated. The HDX-MS data in panel **D** is from our previous study (*28*), and is provided as a comparison to panel B.

There were no significant differences in H/D exchange between free p110γ-p84 and p110γ-p84 in the presence of PM mimic vesicles without lipidated activators. This is consistent with the p110γ-p84 complex being primarily in solution in the absence of either HRas or Gβγ. When HRas was present on membrane surfaces there were multiple regions that showed significant differences compared to membranes alone (Fig. 4A). This included increased exchange in the helical domain, and multiple regions of the regulatory motif in the kinase domain, as well as decreases in exchange in the kα12 membrane binding C-terminal helix and the HRas interface of the RBD. The changes in the helical and kinase domain are consistent with previously observed conformational changes that accompany membrane binding in p110γ (*28, 34, 39*). Intriguingly, there were almost no significant changes in either p110γ or p84 between Gβγ membranes compared to membranes alone, with only one peptide in the kinase domain showing increased exchange (Fig. 4B).

Consistent with the synergistic activation observed in the lipid kinase assays there were significant differences in exchange observed for p110γ-p84 between HRas membranes and HRas/ Gβγ membranes, including decreased exchange at the helical domain and increased exchange in the regulatory motif of the kinase domain (Fig. 4C). These changes in the helical domain were similar, although of a lesser magnitude than those we have observed when examining binding of p110γ-p101 to Gβγ membranes (Fig. 4D) (*28*), indicating that the binding site for Gβγ on p110γ is conserved between the two complexes. There were no significant decreases in exchange in p84 with Gβγ membranes (Fig. 4C), in contrast to the protection observed in the C-terminal domain of p101 (Fig. 4D), which is consistent with p84 lacking a binding site for Gβγ subunits.

### Analysis of Gβγ binding to p110γ and p101

We have previously extensively characterised Gβγ binding of both the p101 and p110γ subunits using HDX-MS, and identified mutations in either the helical domain of p110γ or C-terminal domain of p101 that prevent Gβγ activation (*28*). To provide additional insight into the molecular basis for how p110γ and p101 interact with Gβγ, and why p84 lacks this ability we carried out alphafold-multimer (*38*) modeling of both interfaces (Fig. S5+S6). The search models converged on a consensus orientation of Gβγ interaction with the p101 C-terminal domain (Fig. S5), and a different consensus orientation of Gβγ interaction with the helical domain (Fig. S6), both with predicted alignment scores and per-residue estimate of confidence (pLDDT) scores (*37*) consistent with excellent model accuracy (S5A+B, S6A+B).

These models of Gβγ binding allowed us to make a schematic of how the p110γ-p101 complex is able to bind two Gβγ subunits (Fig. 5A). Critically, the Gγ subunit of Gβγ is geranylated at its C-terminus, and in our models the Gγ C-terminus is oriented in a direction pointed toward the membrane when p110γ is oriented towards its putative membrane interface. Examining these models compared to other Gβγ complexes showed that the same face of the Gβ subunit that binds to the PH domain of G Protein-Coupled Receptor Kinase 2 (*40*) binds to the C-terminal domain of p101 and the helical domain of p110γ (Fig. 5B-D). The p101 interface with Gβγ is primarily composed of two helices that occur between β8 and β9 of the C-terminal domain, along with an extensive interface at residues 816-830. In the helical domain the interface is entirely composed of the N-terminal helix (annotated as hα1). While peptides spanning the helices in p101 are not observed in any of our HDX-MS analysis, for both p110γ-p101 (helical / p101 site) and p110γ-p84 (only helical site) the largest HDX-MS differences with Gβγ binding occurred at either the helical domain (551–557) or the p101 C-terminal domain (816–830) (Fig. 5E+F). These sites are also where we previously designed complex disrupting mutations for both p101 (DQDE817AAA, and RKIL821AAA) and p110γ (RK552DD) (*28*), providing further validation of the putative interface.

**Figure 5.**
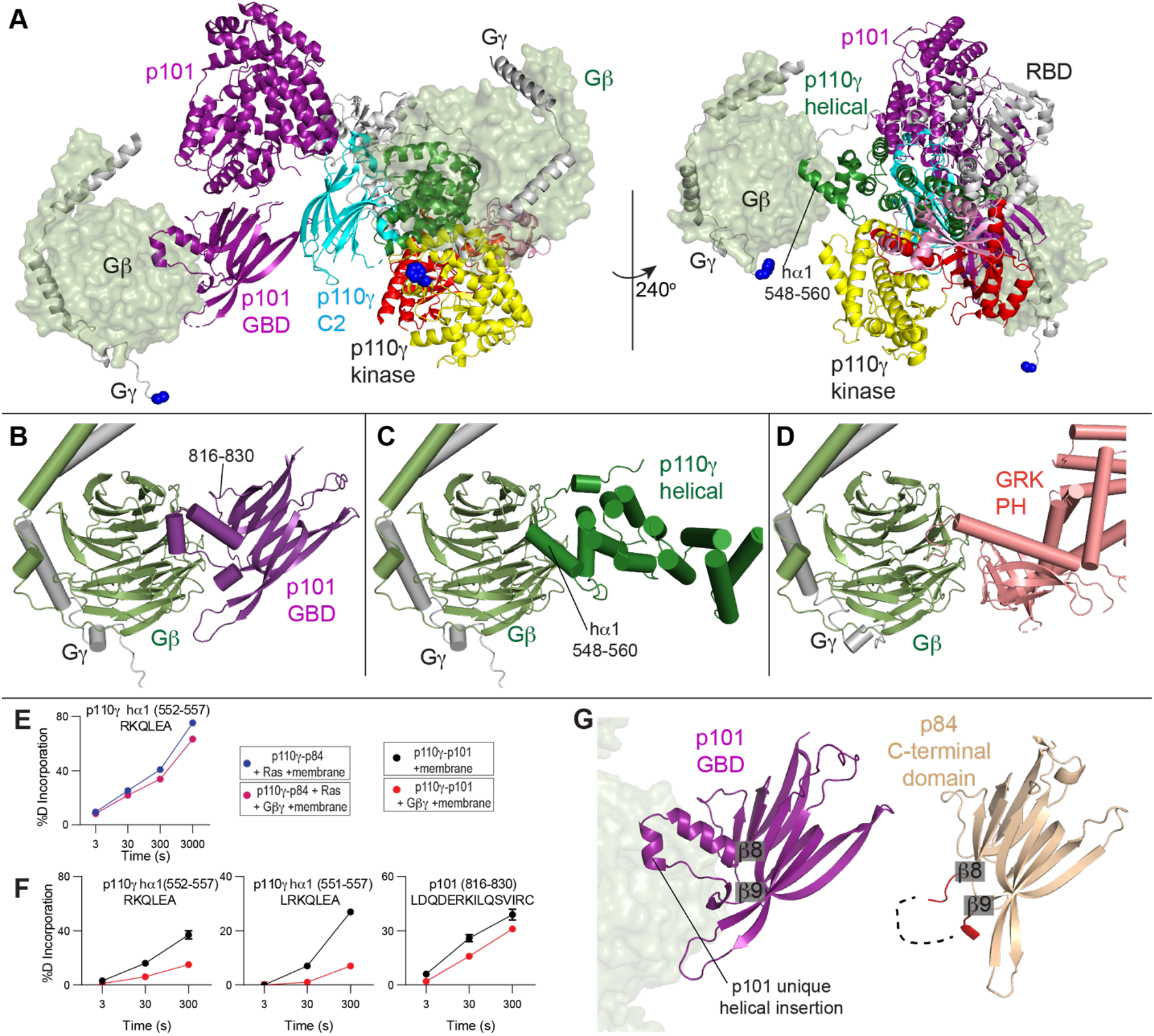
Model of Gβγ activation of PI3Kγ complexes. **A.** Model of the activation of p110γ-p101 complex by two different Gβγ subunits. The location of the Gβγ subunits bound to the C-terminal domain of p101 (Fig. S5) and the helical domain of p110γ (Fig. S6) was based on alphafold2-multimer modelling aligned to the structure of the p110γ-p101 complex (PDB:7MEZ). The domains of p110γ-p101 are annotated, with the Gβ subunit shown as a transparent surface, and the Gγ subunit shown as cartoon, with the C-terminus colored in blue. Both Gβγ subunits are positioned in an orientation compatible with membrane binding of p110γ. **B.** Model of the C-terminal domain of p101 bound to Gβγ (full details on alphafold2-multimer modelling is Fig. S5). The unique helical extension in p101 is annotated, as well as the Gβγ contact surface (816–830). **C.** Model of the helical domain of p110γ bound to Gβγ (full details on alphafold2-multimer modelling is Fig. S6). The N-terminal helix of the helical domain in contact with Gβγ is annotated. **D.** Structure of the PH domain of GPCR kinase 2 (GRK2) bound to Gβγ (PDB: 1OMW) (*40*). **E+F.** Selected deuterium exchange incorporation curves for peptides in the helical domain of p110γ in p110γ-p84 (**E**) and p110γ-p101 (**F**) or C-terminal domain of p101 (**F**) in the presence and absence of Gβγ. Error is shown as standard deviation (n=3). The HDX-MS data in panel **F** is from our previous study (*28*). **G.** Comparison of the C-terminal domain between p101 and p84. The evolutionarily conserved helical extension that occurs between β8 and β9 in p101 is annotated, with the Gβγ subunit from panel B shown as a transparent surface. The end and start of β8 and β9, respectively are labelled, highlighting the corresponding loop between p84 and p101, with the loop coloured red in p84. In p84 the majority of this loop was disordered in both the X-ray and alphafold2-multimer modelling, and is indicated as a dotted line.

There are differences in the orientation and residues mediating Gβγ binding between the p101 and p110γ sites (Fig. 5B+C). The p101 site forms a more extensive interface with Gβγ, with multiple Gβγ contact sites that are unique compared to the helical domain. These differences in interaction are consistent with unique mutations in Gβγ having differential effects between p110γ and p110γ-p101 activation (*36*). Examining the structures of the C-terminal domains showed differences between p101 and p84 at the site where p101 binds Gβγ. Overall, the C-terminal domains are mainly structurally conserved, but the two helices at the interface with Gβγ in p101 between β8 and β9 are absent in p84 (Fig. 5G). This reveals the structural basis for the absence of binding between p84 and Gβγ, and provides a molecular underpinning for the p110γ-p101 complexes sensitivity towards GPCR activation. Overall, this model is consistent with our previous biochemical and TIRF microscopy data supporting the engagement of two Gβγ molecules by the p110γ-p101 complex (*30*).

## Discussion

The class IB PI3Kγ is a key regulator of the immune system (*1, 8, 41*) and is a therapeutic target for multiple human diseases including cancer and inflammatory diseases (*10, 17, 18*). Selective p110γ inhibitors are currently in phase II clinical trials, so fully understanding the regulation of PI3Kγ is essential for continued therapeutic development. The activity of p110γ is fundamentally regulated by its association with either p84 or p101 regulatory subunits, as neutrophils lacking both regulatory subunits have similar PIP_3_ responses to a p110γ kinase dead knock-in mutant (*26*). Here, we report clear molecular insight into how Ras and GPCRs differentially regulate the p110γ-p84 and p110γ-p101 complexes.

The structure of p110γ-p84 reveals that the p84 subunit shares a similar architecture to the p101 subunit (*30*). The p101 and p84 regulatory subunits are differentially expressed in tissues that express p110γ, with biochemical evidence suggesting that p110γ-p84 is dynamic, with p101 able to replace the p84 subunit, and p84 not able to replace p101 (*31*). While the overall secondary structure at the interface with p110γ is conserved, there are numerous evolutionarily conserved differences between p101 and p84 in amino acids at the p110γ interface. We identified two specific cation pi interactions in p110γ-p101, that are absent in p110γ-p84. The dynamic nature of p110γ-p84 has important implications for PI3Kγ signaling and inhibition, as this suggests that any stimuli that may depend on binding or modulating free p110γ will only occur in p110γ-p84. A antibody that bound the p84/p101 interface on the C2 domain of p110γ selectively inhibited only p110γ-p84 and not p110γ-p101. This is likely mediated by p110γ-p84 dissociating, and the antibody sterically preventing regulatory subunit binding, with the antibody binding surface being inaccessible in p110γ-p101 (*42*). The p110γ subunit can be activated by protein kinase C phosphorylation of the helical domain downstream of the IgE antigen receptor in mast cells, with this putatively only occurring for p110γ-p84 and not p110γ-p101 (*43*). This phosphorylation site is in a location that may be inaccessible to p110γ when bound to either p101 or p84, this may provide a unique mechanism for why only p110γ-p84 complexes can be activated by PKC. Further biochemical and structural studies will be required to examine if dynamic differences in p110γ-p84 and p110γ-p101 control regulation by post-translational modifications.

Biochemical assays of HRas activation showed that in the absence of Gβγ both p110γ-p101 and p110γ-p84 are similarly weakly activated by saturating HRas, consistent with previous observations (*28, 29, 31, 32*). Dose response experiments clearly showed that the affinity for HRas activation was equivalent between the two complexes, which is consistent with the Ras interface being distant from the p101-p84 interface (*35*). HDX-MS experiments showed lipidated HRas was able to recruit p110γ-p84 to the membrane, however, it could not fully activate kinase activity. This suggests that HRas by itself acts as a critical regulator of the membrane binding, but both complexes require Gβγ for robust activation. In p110γ-p84 the presence of HRas led to large synergistic activation by Gβγ. This was supported by HDX-MS experiments showing limited Gβγ mediated membrane recruitment of p110γ-p84, and only showed clear differences at the Gβγ interface when both HRas and Gβγ present. For p110γ-p84 with both HRas and Gβγ present at saturating concentrations the kinase activity was still much lower than Gβγ activation of p110γ-p101. This is consistent with the Gβγ interfaces in both the helical domain of p110γ and the GBD of p101 being critical in orienting the p110γ catalytic subunit for maximal kinase activity. This biochemical and biophysical data provides a molecular underpinning for the observation in cells that Ras is required for p110γ-p84 activation (*32*), and for why full activation requires an intact Gβγ biding site in p110γ (*26*). This also explains why in mast cells, which only express p84, inhibitors of Ras lipidation abrogate PI3Kγ signaling, while upon treatment in immune cells expressing p101, PI3Kγ signaling is maintained (*44*).

The p110γ catalytic subunit being almost completely inactive in the absence of a regulatory subunit is unique among class I PI3K isoforms, as in the other class I PI3K isoforms (p110α, p110β, p110δ) the catalytic subunit alone is highly active (*45–47*), and the regulatory subunit acts to inhibit kinase activity and stabilize the catalytic subunit. The p110γ subunit is inhibited through the presence of an internal Tryptophan lock in the regulatory motif of the kinase domain (*34, 39*), with this putatively opened when the p110γ subunit is properly oriented on a membrane surface (*28*). The opening of this lock putatively reorients the C-terminal helix of the kinase domain, allowing it to interact with membrane surfaces, and allowing the activation loop to bind to lipid substrate. The requirement of Gβγ for robust activation of p110γ possibly implies that it orients the catalytic subunit in a manner that disrupts this inhibitory interface. This is supported by cellular experiments that show constitutively membrane localized p110γ is activated by GPCRs (*48*). Additional computational and biophysical studies of p110γ bound to membrane in an inactive and active conformation will be required to fully define the molecular basis for conformational changes required for the fully active state.

Modelling of Gβγ binding to both p101 and p110γ revealed insight into how Gβγ can activate PI3Kγ complexes. This p110γ-p101 can bind two Gβγ subunits, with p110γ-p84 able to bind only a single Gβγ subunit. These models agreed well with our previous HDX-MS and mutational analysis of p101 and p110γ (*28*), as well as TIRF microscopy experiments examining membrane recruitment using varying Gβγ concentrations which implied that the p110γ-p101 complex bound two Gβγ subunits (*30*). The interface in p110γ is located in the N-terminal helix of the helical domain. HDX-MS experiments found this same region mediates Gβγ binding in the class IA PI3K isoform p110β (*49*). Similar to p110γ-p84, p110β requires additional activation and membrane recruitment by either RTKs or Rho GTPases to be robustly activated by Gβγ subunit, suggesting this is either a relatively weak interface, or that binding is dependent on conformational changes induced by membrane binding. Intriguingly, in both p110β and p110γ there is a conformational change in this helix upon membrane recruitment (*28, 49*). The interface in the C-terminal domain of p101 is primarily composed of two helices between β8 and β9, which is evolutionarily conserved in p101, and is not conserved in p84. This provides a molecular underpinning for why p84 shows greatly reduced sensitivity and activation by Gβγ subunits, even in the presence of Ras. The Gβγ interface in p101 is more extensive than that found in p110γ, which may explain why Gβγ alone can so potently activate p110γ-p101, and does not require additional membrane localized activators.

The development of therapeutics targeting PI3Kγ are clinically advanced, with ATP competitive small molecule inhibitors currently in phase II clinical trials in cancer (*19*), and in pre-clinical investigation in chronic obstructive pulmonary disease and inflammatory disease (*12*). There are potential challenges for even highly selective p110γ inhibitors, as immune side effects may be difficult to avoid, highlighted by patients with inactivating primary immunodeficiency clinical p110γ mutations (*20, 21*). The molecular insight into the difference in how p110γ-p101 and p110γ-p84 are regulated could lead to inhibitors specific for either p110γ-p101 or p110γ-p84, which may maintain therapeutic benefit but with decreased side effects. This fits with our observation of nanobodies that block Ras activation strongly inhibit p110γ-p84 activation, while those blocking the p101 interface with Gβγ selectively targeting p110γ-p101 (*50*). Further medicinal chemistry efforts may reveal opportunities to target these sites by small molecule inhibitors.

Collectively, our detailed biochemical and structural analysis of p110γ-p84 and p110γ-p101 provides unique insight into how PI3Kγ complexes are assembled and activated. Our work has defined the molecular basis for how these two distinct complexes can differentially integrate upstream signals, similarly to how different regulatory subunits can alter the activation of mTOR complexes (*51*). A summary of the molecular differences between the p110γ-p84 and p110γ-p101 and their activation by Ras and Gβγ are shown in Fig. 6. This work provides a framework for the design of allosteric modulators for both p110γ-p84 and p110γ-p101, which may inform PI3Kγ complex-specific therapeutic development in inflammatory diseases and cancer.

**Figure 6.**
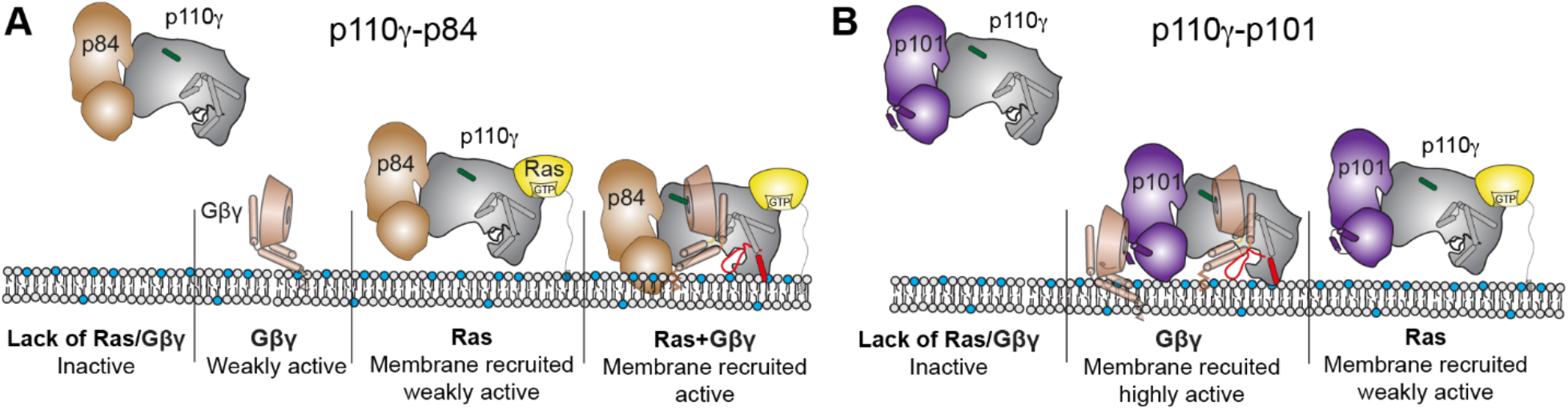
Model of differential activation of PI3Kγ complexes by Gβγ and Ras. **A+B.** Schematic of how Ras and Gβγ subunits can activate p110γ-p84 (**A**) and p110γ-p101 (**B**). Ras in the absence of Gβγ leads to membrane recruitment for both complexes, but only weakly activates kinase activity. The Gβγ binding helices in the GBD of p101 and the helical domain interface with Gβγ are shown, with the helical domain hα1 highlighted in green. The C-terminal helix in the kinase domain that reorients upon membrane binding is highlighted in red upon activation.

## Acknowledgements

We thank Alex Berndt and ESRF ID29 beamline scientist Christoph Müller-Dieckmann for assistance with X-ray diffraction data collection. JEB is supported by an operating grant from the Canadian Institute of Health Research (CIHR, 168998), with salary support from the Michael Smith Foundation for Health Research Scholar award (MSFHR-17686). C.K.Y. is supported by CIHR (FDN-143228) and the Natural Sciences and Engineering Research Council of Canada (RGPIN-2018-03951). RLW is supported by the Medical Research Council (MC_U105184308) and Cancer Research UK (grant DRCPGM\100014). XZ was supported by an MRC-LMB Cambridge Scholarship and by the Cambridge Overseas Trust.

## Conflict of Interest statement

JEB reports personal fees from Scorpion Therapeutics and Olema Oncology; and research grants from Novartis. Other authors declare no competing interests.

## Data Availability

The mass spectrometry proteomics data have been deposited to the ProteomeXchange Consortium via the PRIDE partner repository (*65*). The coordinates for the p110γ-p84 complex have been deposited in the protein data bank with the identifier 8AJ8, and the negative stain EM dataset have been deposited at the EM data bank with the identifier 27738.

## Methods

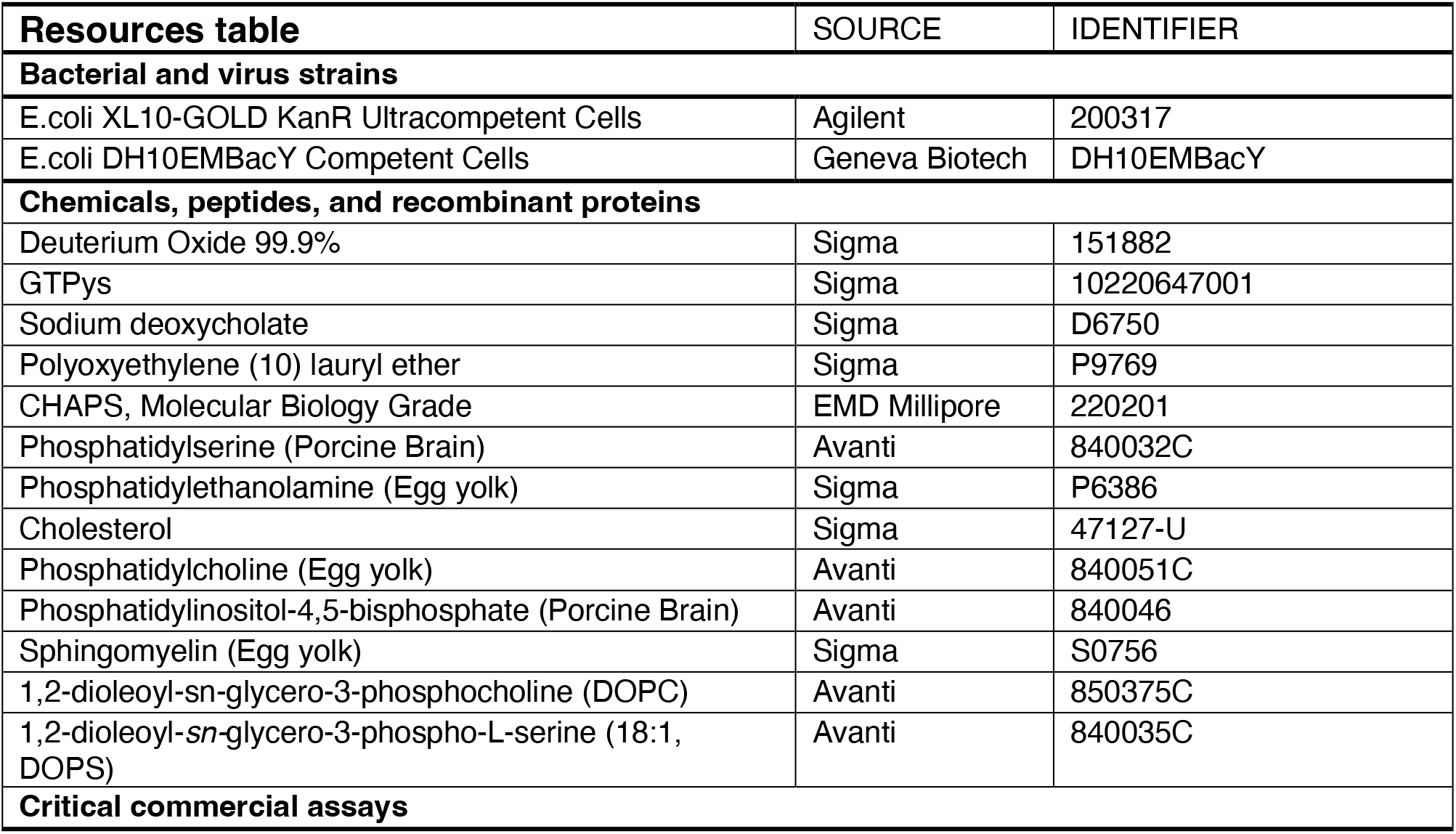
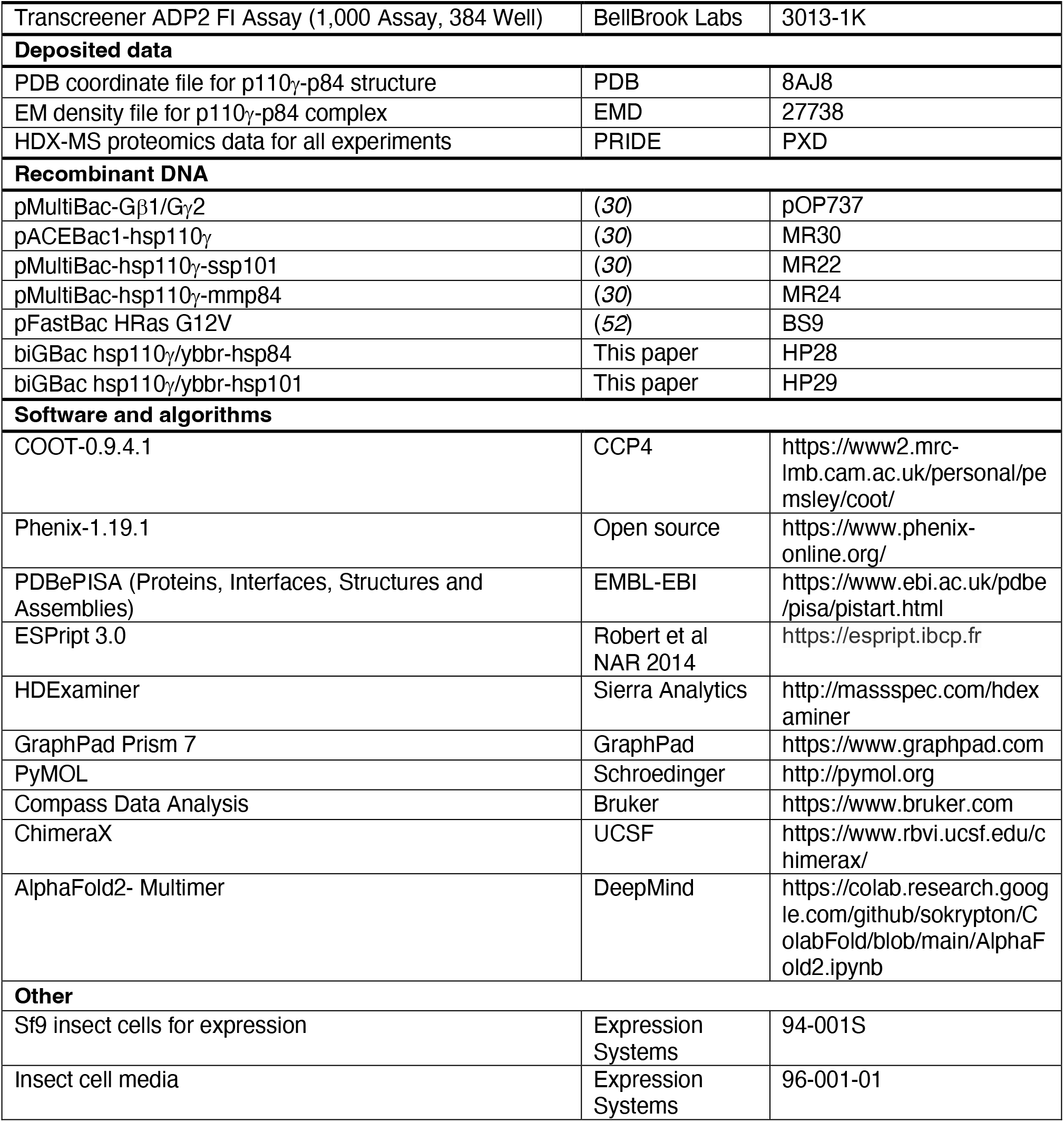

### Plasmid Generation

Plasmids encoding Homo sapiens p110γ (human), Mus musculus p84 (mouse), Sus scrofa p101 (porcine), and Gβγ were used as previously described (*30*). The full-length human PIK3R5 (p101) gene was purchased from Addgene (70464), and the full-length human PIK3R6 (p84) gene was purchased from DanaFarber (HsCD00462228). Plasmids encoding HRas was used as previously described (*52*). PI3K genes were subcloned into pLIB vectors for expression with no engineered tags, while in the case of p110γ a TEV cleavable C-terminal 10x histidine and 2x strep tag was added. Genes were subsequently amplified following the biGBac protocol to generate plasmids containing hsP110γ/hsP101 and hsP110γ/hsP84.

For purification, a 10× histidine tag, a 2× strep tag, and a tobacco etch virus protease cleavage site were cloned to the N terminus of the regulatory subunits for the complex and to p110γ for constructs without regulatory subunits.

### Virus Generation and Amplification

The plasmids encoding genes for insect cell expression were transformed into DH10MultiBac cells (MultiBac, Geneva Biotech) to generate baculovirus plasmid (bacmid) containing the genes of interest. Successful generation was identified by blue-white colony screening and the bacmid was purified using a standard isopropanol-ethanol extraction method. Bacteria were grown overnight (16 hours) in 3-5 mL 2xYT (BioBasic #SD7019). Cells were spun down and the pellet was resuspended in 300 μL of 50 mM Tris-HCl, pH 8.0, 10 mM EDTA, 100 mg/mL RNase A. The pellet was lysed by the addition of 300 μL of 1% sodium dodecyl sulfate (SDS) (W/V), 200 mM NaOH, and the reaction was neutralized by addition of 400 μL of 3.0 M potassium acetate, pH 5.5. Following centrifugation at 21130 RCF and 4 °C (Rotor #5424 R), the supernatant was mixed with 800 μL isopropanol to precipitate bacmid DNA. Following centrifugation, the pelleted bacmid DNA was washed with 500 μL 70% Ethanol three times. The pellet was then air dried for 1 minute and re-suspended in 50 μL Buffer EB (10 mM Tris-Cl, pH 8.5; All buffers from QIAprep Spin Miniprep Kit, Qiagen #27104). Purified bacmid was then transfected into Sf9 cells. 2 mL of Sf9 cells at 0.6X106 cells/mL were aliquoted into a 6-well plate and allowed to attach to form a confluent layer. Transfection reactions were prepared mixing 8-12 μg of bacmid DNA in 100 μL 1xPBS and 12 μg polyethyleneimine (Polyethyleneimine ‘‘Max’’ MW 40.000, Polysciences #24765, USA) in 100 μL 1xPBS and the reaction was allowed to proceed for 20-30 minutes before addition to an Sf9 monolayer containing well. Transfections were allowed to proceed for 5-6 days before harvesting virus containing supernatant as a P1 viral stock.

Viral stocks were further amplified by adding P1 to Sf9 cells at ∼2×10^6^ cells/mL (2/100 volume ratio). This amplification was allowed to proceed for 4-5 days and resulted in a P2 stage viral stock that was used in final protein expression. Harvesting of P2 viral stocks was carried out by centrifuging cell suspensions in 50 mL Falcon tubes at 2281 RCF (Beckman GS-15). To the supernatant containing virus, 5-10% inactivated fetal bovine serum (FBS; VWR Canada #97068-085) was added and the stock was stored at 4°C.

### Expression and purification of PI3Kγ constructs

All PI3K*γ* constructs were expressed in Sf9 insect cells using the baculovirus expression system. Following 55 hours of expression, cells were harvested by centrifuging at 1680 RCF (Eppendorf Centrifuge 5810 R) and the pellets were snap-frozen in liquid nitrogen. The complex was purified through a combination of nickel affinity, streptavidin affinity and size exclusion chromatographic techniques.

Frozen insect cell pellets were resuspended in lysis buffer (20 mM Tris pH 8.0, 100 mM NaCl, 10 mM imidazole pH 8.0, 5% glycerol (v/v), 2 mM βME), protease inhibitor (Protease Inhibitor Cocktail Set III, Sigma)) and sonicated for 2 minutes (15s on, 15s off, level 4.0, Misonix sonicator 3000). Triton-X was added to the lysate to a final concentration of 0.1% and clarified by spinning at 15,000 RCF at 4°C for 45 minutes (Beckman Coulter JA-20 rotor). The supernatant was loaded onto a 5 mL HisTrap™ FF crude column (GE Healthcare) equilibrated in NiNTA A buffer (20 mM Tris pH 8.0, 100 mM NaCl, 20 mM imidazole pH 8.0, 5% (v/v) glycerol, 2 mM βME). The column was washed with high salt NiNTA A buffer (20 mM Tris pH 8.0, 1 M NaCl, 20 mM imidazole pH 8.0, 5% (v/v) glycerol, 2 mM βME), NiNTA A buffer, 6% NiNTA B buffer (20 mM Tris pH 8.0, 100 mM NaCl, 250 mM imidazole pH 8.0, 5% (v/v) glycerol, 2 mM βME) and the protein was eluted with 100% NiNTA B. The eluent was loaded onto a 5 mL StrepTrap™ HP column (GE Healthcare) equilibrated in gel filtration buffer (20mM Tris pH 8.5, 100 mM NaCl, 50 mM Ammonium Sulfate and 0.5 mM TCEP). The column was washed with the same buffer and loaded with tobacco etch virus protease. After cleavage on the column overnight, the protein was eluted in gel filtration buffer. The protein was concentrated in a 50,000 MWCO Amicon Concentrator (Millipore) to <1 mL and injected onto a Superdex™ 200 10/300 GL Increase size-exclusion column (GE Healthcare) equilibrated in gel filtration buffer. After size exclusion, the protein was concentrated, aliquoted, frozen, and stored at −80°C.

### Expression and Purification of lipidated Gβγ for kinase activity assays

Full length, lipidated human Gβγ (Gβ1γ2) was expressed in Sf9 insect cells and purified as described previously (*53*). After 65 hours of expression, cells were harvested, and the pellets were frozen as described above. Pellets were resuspended in lysis buffer (20 mM HEPES pH 7.7, 100 mM NaCl, 10 mM βME, protease inhibitor (Protease Inhibitor Cocktail Set III, Sigma)) and sonicated for 2 minutes (15s on, 15s off, level 4.0, Misonix sonicator 3000). The lysate was spun at 500 RCF (Eppendorf Centrifuge 5810 R) to remove intact cells and the supernatant was centrifuged again at 25,000 RCF for 1 hour (Beckman Coulter JA-20 rotor). The pellet was resuspended in lysis buffer and sodium cholate was added to a final concentration of 1% and stirred at 4°C for 1 hour. The membrane extract was clarified by spinning at 10,000 RCF for 30 minutes (Beckman Coulter JA-20 rotor). The supernatant was diluted 3 times with NiNTA A buffer (20 mM HEPES pH 7.7, 100 mM NaCl, 10 mM Imidazole, 0.1% C12E10, 10mM βME) and loaded onto a 5 mL HisTrap™ FF crude column (GE Healthcare) equilibrated in the same buffer. The column was washed with NiNTA A, 6% NiNTA B buffer (20 mM HEPES pH 7.7, 25 mM NaCl, 250 mM imidazole pH 8.0, 0.1% C12E10, 10 mM βME) and the protein was eluted with 100% NiNTA B. The eluent was loaded onto HiTrap™ Q HP anion exchange column equilibrated in Hep A buffer (20 mM Tris pH 8.0, 8 mM CHAPS, 2 mM Dithiothreitol (DTT)). A gradient was started with Hep B buffer (20 mM Tris pH 8.0, 500 mM NaCl, 8 mM CHAPS, 2 mM DTT) and the protein was eluted in ∼50% Hep B buffer. The eluent was concentrated in a 30,000 MWCO Amicon Concentrator (Millipore) to < 1 mL and injected onto a Superdex™ 75 10/300 GL size exclusion column (GE Healthcare) equilibrated in Gel Filtration buffer (20 mM HEPES pH 7.7, 100 mM NaCl, 10 mM CHAPS, 2 mM TCEP). Fractions containing protein were pooled, concentrated, aliquoted, frozen and stored at −80 °C.

### Expression and Purification of Lipidated HRas G12V

Full-length HRas G12V was expressed by infecting 500 mL of Sf9 cells with 5 mL of baculovirus. Cells were harvested after 55 hours of infection and frozen as described above. The frozen cell pellet was resuspended in lysis buffer (50 mM HEPES pH 7.5, 100 mM NaCl, 10 mM βME and protease inhibitor (Protease Inhibitor Cocktail Set III, Sigma)) and sonicated on ice for 1 minute 30 seconds (15s ON, 15s OFF, power level 4.0) on a Misonix sonicator 3000. Triton-X 114 was added to the lysate to a final concentration of 1%, mixed for 10 minutes at 4°C and centrifuged at 25,000 rpm for 45 minutes (Beckman Ti-45 rotor). The supernatant was warmed to 37°C for few minutes until it turned cloudy following which it was centrifuged at 11,000 rpm at room temperature for 10 minutes (Beckman JA-20 rotor) to separate the soluble and detergent-enriched phases. The soluble phase was removed, and Triton-X 114 was added to the detergent-enriched phase to a final concentration of 1%. Phase separation was performed 3 times. Imidazole pH 8.0 was added to the detergent phase to a final concentration of 15 mM and the mixture was incubated with Ni-NTA agarose beads (Qiagen) for 1 hour at 4°C. The beads were washed with 5 column volumes of Ras-NiNTA buffer A (20mM Tris pH 8.0, 100mM NaCl, 15mM imidazole pH 8.0, 10mM βME and 0.5% Sodium Cholate) and the protein was eluted with 2 column volumes of Ras-NiNTA buffer B (20mM Tris pH 8.0, 100mM NaCl, 250mM imidazole pH 8.0, 10mM βME and 0.5% Sodium Cholate). The protein was buffer exchanged to Ras-NiNTA buffer A using a 10,000 kDa MWCO Amicon concentrator, where protein was concentrated to ∼1mL and topped up to 15 mL with Ras-NiNTA buffer A and this was repeated a total of 3 times. GTPγS was added in 2-fold molar excess relative to HRas along with 25 mM EDTA. After incubating for an hour at room temperature, the protein was buffer exchanged with phosphatase buffer (32 mM Tris pH 8.0, 200 mM Ammonium Sulphate, 0.1 mM ZnCl2, 10 mM βME and 0.5% Sodium Cholate). 1 unit of immobilized calf alkaline phosphatase (Sigma) was added per milligram of HRas along with 2-fold excess nucleotide and the mixture was incubated for 1 hour on ice. MgCl2 was added to a final concentration of 30 mM to lock the bound nucleotide. The immobilized phosphatase was removed using a 0.22-micron spin filter (EMD Millipore). The protein was concentrated to less than 1 mL and was injected onto a Superdex 75 10/300 GL size exclusion column (GE Healthcare) equilibrated in gel filtration buffer (20 mM HEPES pH 7.7, 100 mM NaCl, 10 mM CHAPS, 1 mM MgCl2 and 2 mM TCEP). The protein was concentrated to 1 mg/mL using a 10,000 kDa MWCO Amicon concentrator, aliquoted, snap-frozen in liquid nitrogen and stored at −80°C.

### Expression and purification of complex of porcine p110γ with mouse p84

Constructs of full-length porcine p110*γ* were cloned into pVL1393 (Invitrogen). The plasmid for EE-tagged mouse p84 was a gift from Len Stephens (The Babraham Institute, UK). The constructs were transfected into *Spodoptera frugiperda* 9 (Sf9) insect cells with ExGen500 (Fermentas) and incubated at 27°C for 5 days to make baculoviruses. The heterodimeric p110γ-p84 complexes were obtained by co-infection of 3 L of SF9 cells with p110γ-expressing and p84-expressing viruses. Cells were inoculated at a density of 1×10^6^ cells/ml and grown in 2L roller bottles standing vertically, with 500 ml of Sf9 cells per bottle. After 62 hours incubation at 27°C, cells were harvested, washed in PBS, pelleted, snap-frozen in liquid nitrogen and stored at −80°C.

### Purification of complex of porcine p110γ with mouse p84

Frozen cells were resuspended in sonication buffer (50mM TrisHCl pH 8, 100 mM NaCl, 1 mM PEFA, 25 mM imidazole) and lysed by sonication on ice at power 8 for 10 minutes (Sf9 cells). The lysates were ultracentrifuged at 35,000 rpm for 45 minutes at 4°C in Ti45 rotor. The soluble cell lysate was filtered through a 0.45 µm filter. Subsequently, the lysate was passed over a 5 ml Ni-NTA Fast Flow column (GE Healthcare) that had been equilibrated with Ni wash buffer (20 mM Tris pH8, 1% Betaine, 0.1 M NaCl, 50 mM potassium phosphate pH 7, 0.05% Tween), washed with 15 ml of Ni wash buffer then eluted in a gradient from Ni A buffer (20 mM Tris pH 8, 300 mM NaCl, 25 mM imidazole) to Ni B (20 mM Tris pH 8, 300 mM NaCl, 500 mM imidazole). Fractions containing the p110-p84 complex were pooled and diluted 1:2 with QA buffer (50 mM Tris pH 8, 2 mM DTT). The diluted sample was loaded onto tandem HiTrap Q (5ml, GE Healthcare) and HiTrap Heparin (5 ml, GE Healthcare) columns that had been equilibrated in tandem with QA buffer. The protein was eluted from the tandem columns with a gradient of QA buffer to QB buffer (50 mM Tris pH 8, 1 M NaCl, 2 mM DTT). The eluted fractions containing the heterodimer were pooled and concentrated to 2 mL in a 50 kD MWCO Amicon Ultra concentrator (Millipore). The concentrated sample was then purified using a Superdex 200 (16/60) gel-filtration column with gel filtration buffer (20 mM Tris pH 7.5, 100 mM NaCl, 2 mM DTT). The fractions containing the heterodimer were pooled and concentrated to 10 mg/ml. One preparation from 3L of Sf9 cells yielded about 11 mg of purified, concentrated heterodimer.

### Alphafold2 modelling

We utilized the AlphaFold2 using MMseqs2 notebook of ColabFold at colab.research.google.com/github/sokrypton/ColabFold/blob/main/AlphaFold2.ipynb (*54*) to make structural predictions of p110γ bound to p84, p101 bound to Gβγ, and p110γ bound to Gβγ. The pLDDT confidence values consistently scored above 90% for all models, with the predicted aligned error and pLDDT scores for all models are shown in Figs. S2, S5, S6. The best models for Gβγ bound to the helical domain of p110γ and the C-terminal domain of p101 are included as PDB files in the source data.

### Crystallization of porcine p110γ/mouse p84

For initial screens, 100 nL drops of purified, concentrated heterodimer at 10 mg/ml were dispensed into LMB 96-well plates with 100 nL of reservoir solution. The initial screen was the 2000 condition LMB screen (*55, 56*), containing a wide range of crystallisation solutions, using an Innovadyne crystallisation robot. The plates were stored at 17°C. To improve initial crystals, 1μl drops of protein and 1μl drops of well solution were manually pipetted into 24 well plates (either sitting drop or hanging drop). Seeding from the existing crystals into the fresh drop was performed using a Hampton seeding tool. The plates were then stored at 17°C or 4°C. Crystals were initially obtained from a Morpheus screen (*57*). Optimized crystals were grown from a crystallization solution containing 16% EDO_P8K (20% w/v PEG 8000, 40% v/v ethylene glycol), 0.06 M amino acids (0.2 M sodium L-glutamate, 0.2 M DL-alanine, 0.2 M glycine, 0.2 M DL-lysine, 0.2 M DL-serine), 0.08 M buffer 2 pH7.5 (0.5 M HEPES, 0.5 M MOPS), 0.4 M Na/K phosphate pH 6.3. Crystals were 120μm x 50μm x 10μm plates that diffracted to 8 Å resolution (Table S1).

### X-ray data collection/refinement for complex of porcine p110γ with mouse p84

Diffraction data collected with remote control at ESRF beamline ID29, using a wavelength of 0.9762. Images were integrated with MOSFLM (*58*) and scaled with SCALA (*59*). Molecular replacement and refinement were carried out using PHASER (*60*) and Phenix.refine (*61*). For molecular replacement a model of the p110γ-p84 complex was generated in COOT (*62*) from a composite of the alphafold2 model of the p110γ C2 domain and RBD-C2 and C2-helical linkers bound to full length p84 with the rest of the sus scrofus p110γ subunit assembled from an alphafold generated model templated on the human p110γ from the PDB entry 7MEZ (*30*). There were four heterodimers per asymmetric unit. The entire assembly was then subjected to rigid-body, xyz reciprocal space, and group B-factor refinement in phenix-refine (*61*) using NCS and secondary structure restraints. Due to the low resolution, no manual adjustments were made in the model. Statistics for the final model are shown in Table S1.

### Negative stain electron microscopy

Purified p110γ-mmp84 was adsorbed to glow discharged carbon coated grids at a concentration of 0.02 mg/mL for 5s and stained with uranyl formate. The stained specimen was examined using a Tecnai Spirit transmission electron microscope (ThermoFisher Scientific) operated at an accelerating voltage of 120 kV and equipped with an FEI Eagle 4K charged-coupled-device (CCD) camera. 50 micrographs were acquired at a nominal magnification of 49,000x at a defocus of −1.2mm and binned twice to obtain a final pixel size of 4.67 Å/pixel. The contrast transfer function (CTF) of each micrograph was estimated using CTFFind4.1 within Relion 3.0.8. 200 particles were manually picked then aligned to generate 2D class averages for template-based autopicking. These templates were then used to autopick 20,610 particles which were extracted with a box size of 336 Å. Particles were then exported to cryoSPARC v2.14.2 for 2D classification and 10,344 particles which classified to “good” classes were selected and subjected to ab initio reconstruction with a max alignment resolution of 12 Å. The same particles were then used for homogenous refinement of the *ab initio* model, yielding the final map at 19 Å resolution, as calculated by the gold standard Fourier Shell Correlation (FSC) at 0.143 cutoff.

### Lipid vesicle preparation for kinase activity assays

Lipid vesicles containing 5% brain phosphatidylinositol 4,5-bisphosphate (PIP2), 20% brain phosphatidylserine (PS), 35% egg-yolk phosphatidylethanolamine (PE), 10% egg-yolk phosphatidylcholine (PC), 25% cholesterol and 5% egg-yolk sphingomyelin (SM) were prepared by mixing the lipids solutions in organic solvent. The solvent was evaporated in a stream of argon following which the lipid film was desiccated in a vacuum for 45 minutes. The lipids were resuspended in lipid buffer (20 mM HEPES pH 7.0, 100 mM NaCl and 10 % glycerol) at a concentration of 5 mg/ml and the solution was bath sonicated for 15 minutes. The vesicles were subjected to five freeze thaw cycles and extruded 11 times through a 100-nm filter (T&T Scientific: TT-002-0010). The extruded vesicles were aliquoted and stored at −80°C.

### Kinase Assays

All kinase assays were done using Transcreener ADP2 Fluorescence Intensity (FI) assays (Bellbrook labs) which measures ADP production. PM-mimic vesicles [5% phosphatidylinositol 4,5-bisphosphate (PI(4, 5) P2), 20% phosphatidylserine (PS), 10% phosphatidylcholine (PC), 35% phosphatidylethanolamine (PE), 25% cholesterol, 5% sphingomyelin (SM)] at final concentration of 0.5 mg/mL, ATP at a final concentration of 100 μM and HRas at final concentrations ranging from 10 nM to 1.5 μM were used. For assays measuring co-stimulation with Gβγ, 1.5 μM of the activator was used in the reaction. Final concentration of kinase ranged from 400 nM to 2000 nM for both p110γ-p101 and p110γ-p84. For conditions with Gβγ, final kinase concentrations of kinase ranged from 100 nM to 400 nM for p110γ-p84 and from 3 nM to 10 nM for p110γ-p101.

2 μL of 2X substrate solution containing vesicles, the appropriate concentration of Ras and Gβγ (for conditions assaying co-stimulation) was mixed with 2 μL of 2X kinase solution and the reaction was allowed to proceed for 60 minutes. The reactions were stopped with 4 μL of 2X stop and detect solution containing Stop and Detect buffer, 8 nM ADP Alexa Fluor 594 Tracer and 93.7 μg/mL ADP2 Antibody IRDye QC-1 and incubated for 50 minutes. The fluorescence intensity was measured using a SpectraMax M5 plate reader at excitation 590 nm and emission 620 nm. The % ATP turnover was interpolated from a standard curve (0.1-100 μM ADP) using Graphpad prism, with these values converted into specific activity based on the concentration of protein.

### Hydrogen Deuterium eXchange Mass Spectrometry-Activators

Exchange reactions were carried out at 18°C in 12 µL volumes with final concentrations of 1.5 µM, 3 µM, 3µM for p110*γ*-p84, HRas (G12V) and Gβγ respectively. A total of five conditions were assessed: p110γ-p84, p110γ-p84 + HRas (G12V), p110γ-p84 + Gβγ, and p110γ-p84 + HRas (G12V) + Gβγ. All conditions were in the presence of PM mimic membranes [5% phosphatidylinositol 4,5-bisphosphate (PI(4, 5)P2), 20% phosphatidylserine (PS), 10% phosphatidylcholine (PC), 35% phosphatidylethanolamine (PE), 25% cholesterol, 5% sphingomyelin (SM)] at a final concentration of 0.42 mg/ml. Mixtures of lipid vesicles and activators (HRas(G12V)/Gβγ) were prepared by combining 1 µL of lipid vesicles or vesicle buffer (25mM HEPES 7.0, 100mM NaCl, 10% glycerol) with 0.85 µL of HRas(G12V) or HRas buffer (20mM HEPES pH 7.7, 100mM NaCl, 10mM CHAPS, 2mM TCEP), and 0.63 µL of Gβγ or Gβγ buffer (20mM HEPES pH 7.7, 100mM NaCl, 8mM CHAPS, 2mM TCEP). Prior to the addition of D_2_O, 1.2 µL of p110γ-p84 was added to the lipid-activator mixture, and the solution was left to incubate at 18°C for 2 mins. The hydrogen-deuterium exchange reaction was initiated by the addition of 8.32 µL D_2_O buffer (94.3% D_2_O, 100 mM NaCl, 20 mM HEPES pH 7.5) to the 3.68 µL protein or protein-lipid solutions for a final D_2_O concentration of 65.5%. Exchange was carried out over four time points (3s, 30s, 300s, 3000s) and terminated by the addition of 60 µL ice-cold acidic quench buffer (0.6 M guanidine-HCl, 0.9% formic acid final).

### Hydrogen Deuterium eXchange Mass Spectrometry-Regulators

Exchange reactions were carried out at 18°C in 50 µL volumes with final concentrations of 0.4 µM, human p110*γ*/mouse p84 or human p110*γ*/porcine p101. The hydrogen-deuterium exchange reaction was initiated by the addition of 48.6 µL D_2_O buffer (94.3% D_2_O, 100 mM NaCl, 20 mM HEPES pH 7.5) to the 1.4 µL protein solutions for a final D_2_O concentration of 91.7% Exchange was carried out over five time points (3s, 30s, 300s, 3000s at 18°C and 3s at 4°C) and terminated by the addition of 20 µL ice-cold acidic quench buffer (0.6 M guanidine-HCl, 0.9% formic acid final).

### Hydrogen Deuterium eXchange Mass Spectrometry-human regulators

Exchange reactions were carried out at 18°C in either 6ul (high concentration) or 50 µL(low concentration) volumes with final concentrations of 1.5 µM(high) or 0.175uM (low) human p110*γ*/mouse p84 or human p110*γ*/porcine p101. The hydrogen-deuterium exchange reaction was initiated by the addition of 3 µL or 25 µL D_2_O buffer (94.3% D_2_O, 100 mM NaCl, 20 mM HEPES pH 7.5) to the 3µL or 25 µL protein solutions for a final D_2_O concentration of 47.2% Exchange was carried out over two time points (30s, 300s at 18°C) and terminated by the addition of 64 µL or 20 µL ice-cold acidic quench buffer (0.6 M guanidine-HCl, 0.9% formic acid final).

### Protein Digestion and MS/MS Data Collection

Protein samples were rapidly thawed and injected onto an integrated fluidics system containing a HDx-3 PAL liquid handling robot and climate-controlled chromatography system (LEAP Technologies), a Dionex Ultimate 3000 UHPLC system, as well as an Impact HD QTOF Mass spectrometer (Bruker). The protein was run over two immobilized pepsin columns (Applied Biosystems; Poroszyme™ Immobilized Pepsin Cartridge, 2.1 mm x 30 mm; Thermo-Fisher 2-3131-00; at 10°C and 2°C respectively),or for the high low human regulator HDX over one immobilized Nepenthesin-2 column from Affipro (AP-PC-004), at 200 μL/min for 3 minutes. The resulting peptides were collected and desalted on a C18 trap column [Acquity UPLC BEH C18 1.7 mm column (2.1 x 5 mm); Waters 186003975]. The trap was subsequently eluted in line with an ACQUITY 1.7 µm particle, 100 x 1 mm2 C18 UPLC column (Waters 186002352), using a gradient of 3-35% B (buffer A, 0.1% formic acid; buffer B, 100% acetonitrile) over 11 min immediately followed by a gradient of 35-80% B over 5 minutes. MS experiments acquired over a mass range from 150 to 2200 mass/charge ratio (m/z) using an electrospray ionization source operated at a temperature of 200°C and a spray voltage of 4.5 kV.

### Peptide Identification

Peptides were identified using data-dependent acquisition following tandem MS/MS experiments (0.5 s precursor scan from 150-2000 m/z; twelve 0.25 s fragment scans from 150-2000 m/z). MS/MS datasets were analyzed using PEAKS7 (PEAKS), and a false discovery rate was set at 0.1% using a database of purified proteins and known contaminants (*63*). The search parameters were set with a precursor tolerance of 20 parts per million, fragment mass error 0.02 Da, and charge states from 1 to 8.

### Mass Analysis of Peptide Centroids and Measurement of Deuterium Incorporation

HD-Examiner Software (Sierra Analytics) was used to automatically calculate the level of deuterium incorporation into each peptide. All peptides were manually inspected for correct charge state, correct retention time, and appropriate selection of isotopic distribution. Deuteration levels were calculated using the centroid of the experimental isotope clusters. HDX-MS results are presented with no correction for back exchange shown in the Source data, with the only correction being applied correcting for the deuterium oxide percentage of the buffer used in the exchange. Changes in any peptide at any time point greater than specified cut-offs (5% and 0.3 Da, or 7% and 0.5Da for human regulator HDX) and with an unpaired, two-tailed t-test value of p<0.01 was considered significant.

The raw peptide deuterium incorporation graphs for a selection of peptides with significant differences are shown, with the raw data for all analyzed peptides in the source data. To allow for visualization of differences across all peptides, we utilized number of deuteron difference (#D) plots. These plots show the total difference in deuterium incorporation over the entire H/D exchange time course, with each point indicating a single peptide. These graphs are calculated by summing the differences at every time point for each peptide and propagating the error (example Fig 2E, 4A-C). For a selection of peptides we are showing the %D incorporation over a time course, which allows for comparison of multiple conditions at the same time for a given region (Fig. 5E+F). Samples were only compared within a single experiment and were never compared to experiments completed at a different time with a different final D_2_O level. The data analysis statistics for all HDX-MS experiments are in Table S2 according to the guidelines of (*64*). The mass spectrometry proteomics data have been deposited to the ProteomeXchange Consortium via the PRIDE partner repository (*65*).

## Supplementary Figures + Tables

**Fig S1.**
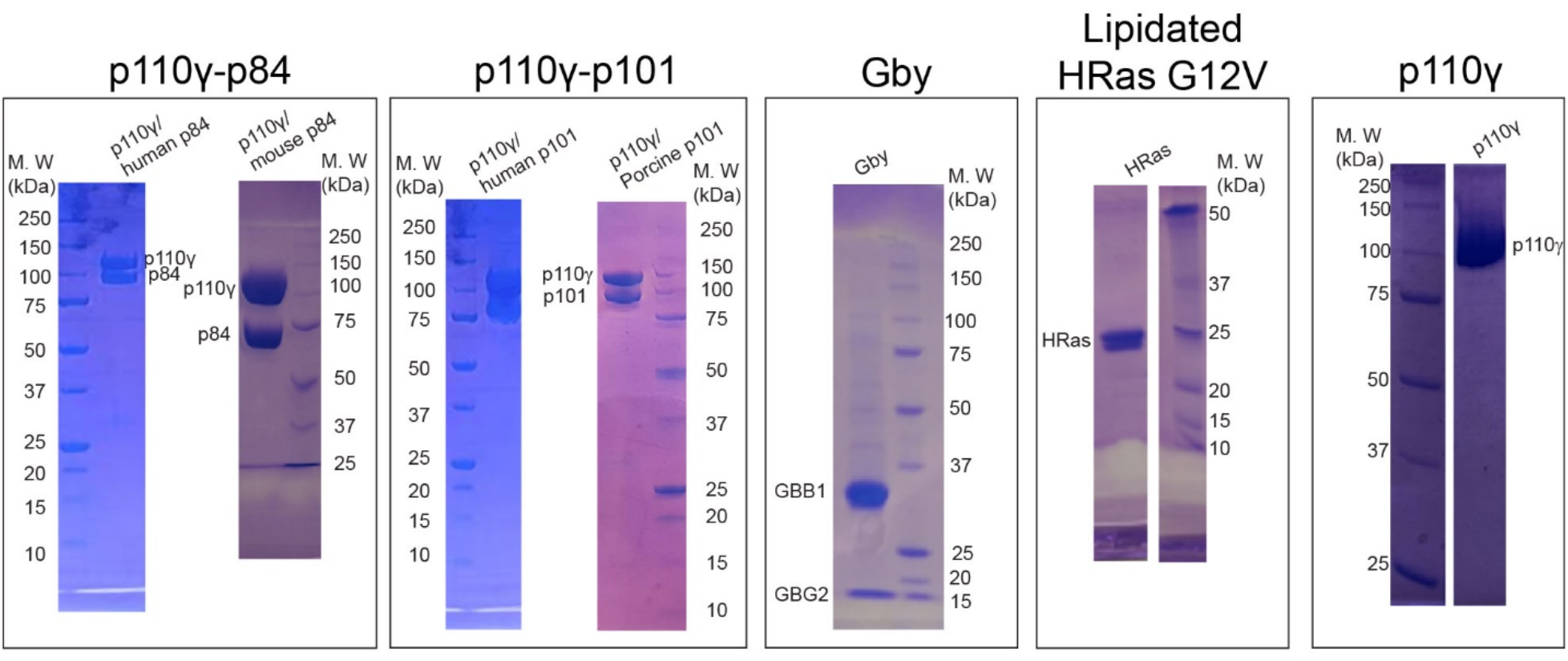
SDS PAGE gel images of all protein constructs used in this study. Full sized images are available in the source data.

**Fig S2.**
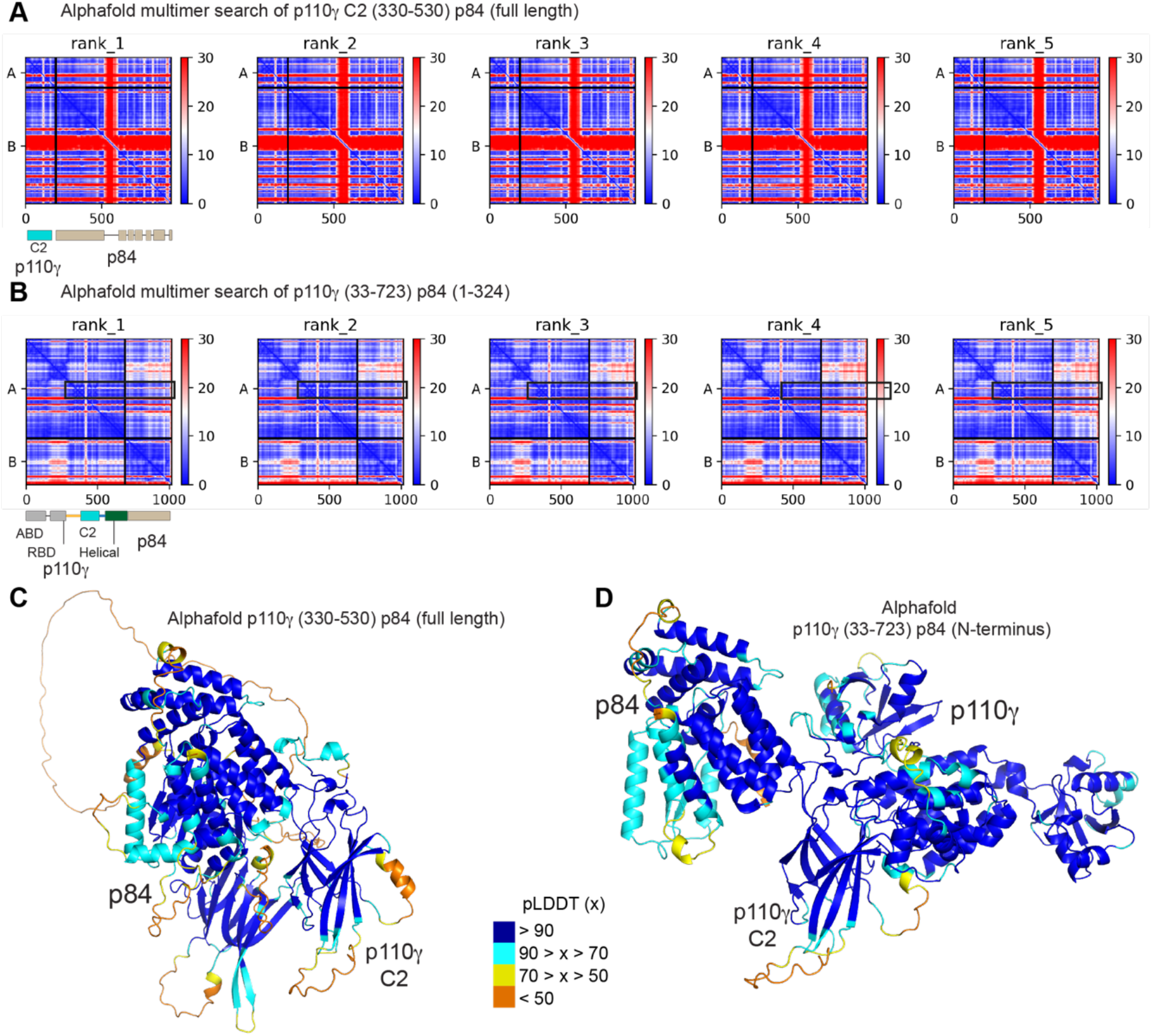
Alphafold2-multimer modelling of the p110γ-p84 complex. **A.** Predicted aligned error (PAE) for Alphafold2 Multimer search of the p110γ C2 domain and RBD-C2 and C2-helical linkers bound to full length p84. The sequence of the two searches are indicated, with a schematic indicated below. **B.** Predicted aligned error (PAE) for Alphafold2 Multimer search of the N-terminus of p110γ (33-723 covering the ABD, RBD, C2, and helical domains) and the N-terminus (1–324) of p84. The sequence of the two searches are indicated, with a schematic indicated below. For both panels **A+B**, the colours indicate the predicted aligned error, and are coloured according to the legend. Note that the PAE plot is not an inter-residue distance map or a contact map. Instead, the red-blue colour indicates expected distance error. The colour at (x, y) corresponds to the expected distance error in residue x’s position, when the prediction are aligned on residue y (more information can be found at https://alphafold.ebi.ac.uk/) (*37, 66*). Blue is indicative of low PAE, with the low PAE at the p110γ-p84 interface in both panels **A+B** suggests that AlphaFold2-multimer predicts the relative positions of the catalytic and regulatory subunits with high accuracy. **C+D.** Alphafold2 models from panels **A+B** shown with the per-residue confidence metric predicted local-distance difference test (pLDDT) coloured according to the legend. The pLDDT score varies from 0 to 100, and is an estimate of how well the prediction would agree with an experimental structure based on the local distance difference test Cα (*37*).

**Fig S3.**
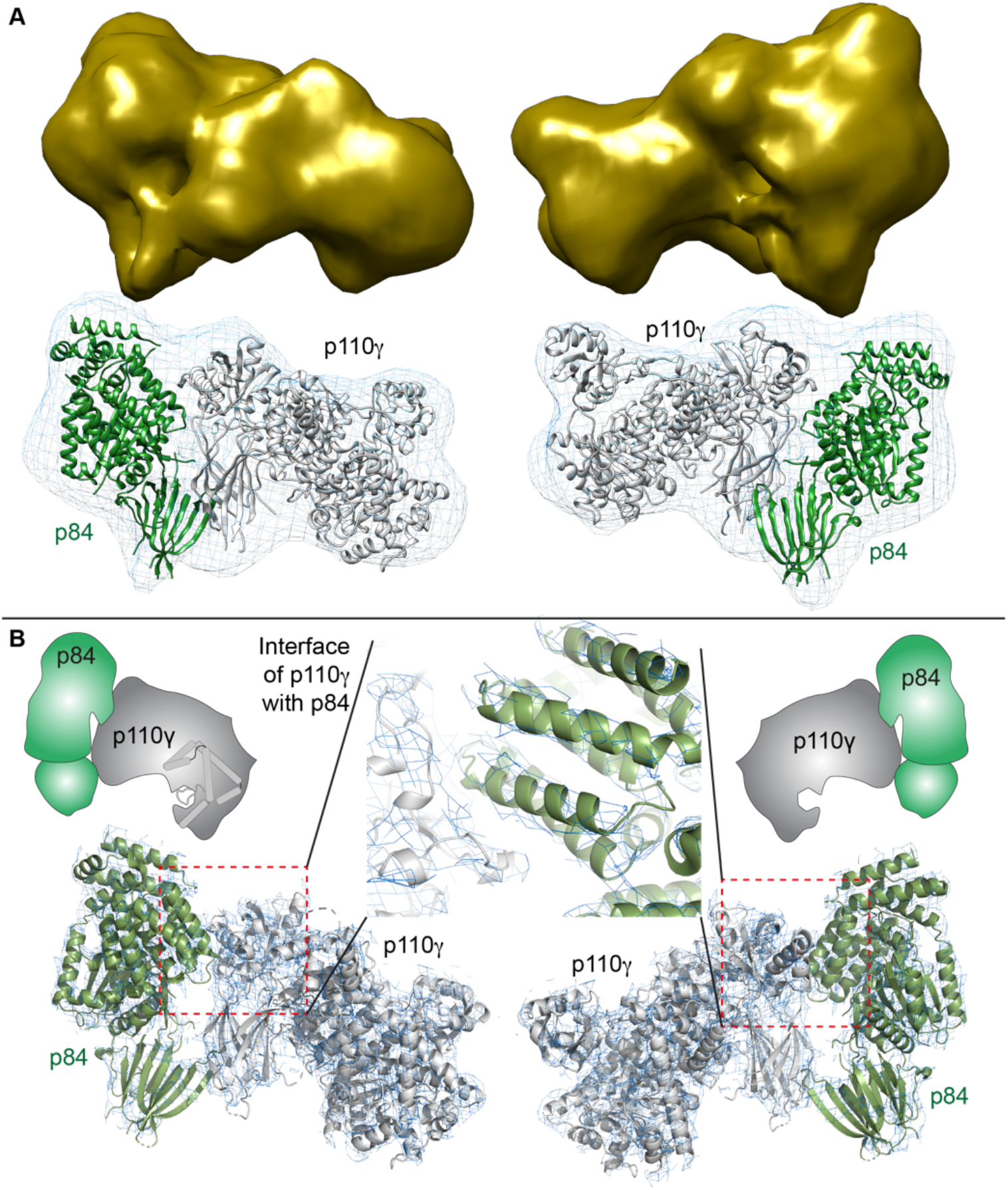
EM and X-ray validation of the alphafold2-multimer model of the p110γ-p84 complex. **A.** 3D EM reconstruction of p110γ with different orientations of the complex. A cartoon representation of the p110y-p84 complex is shown in the density map in the same orientation as above. **B.** The 2Fo-Fc electron density (contoured at 1.5σ) for the complex of the porcine p110γ and mouse p84 complex phased using the alphafold2 generated model. A zoom in of the interface of the p110y-p84 complex shows clear density across for the interfacial helices in p84 (green).

**Fig S4.**
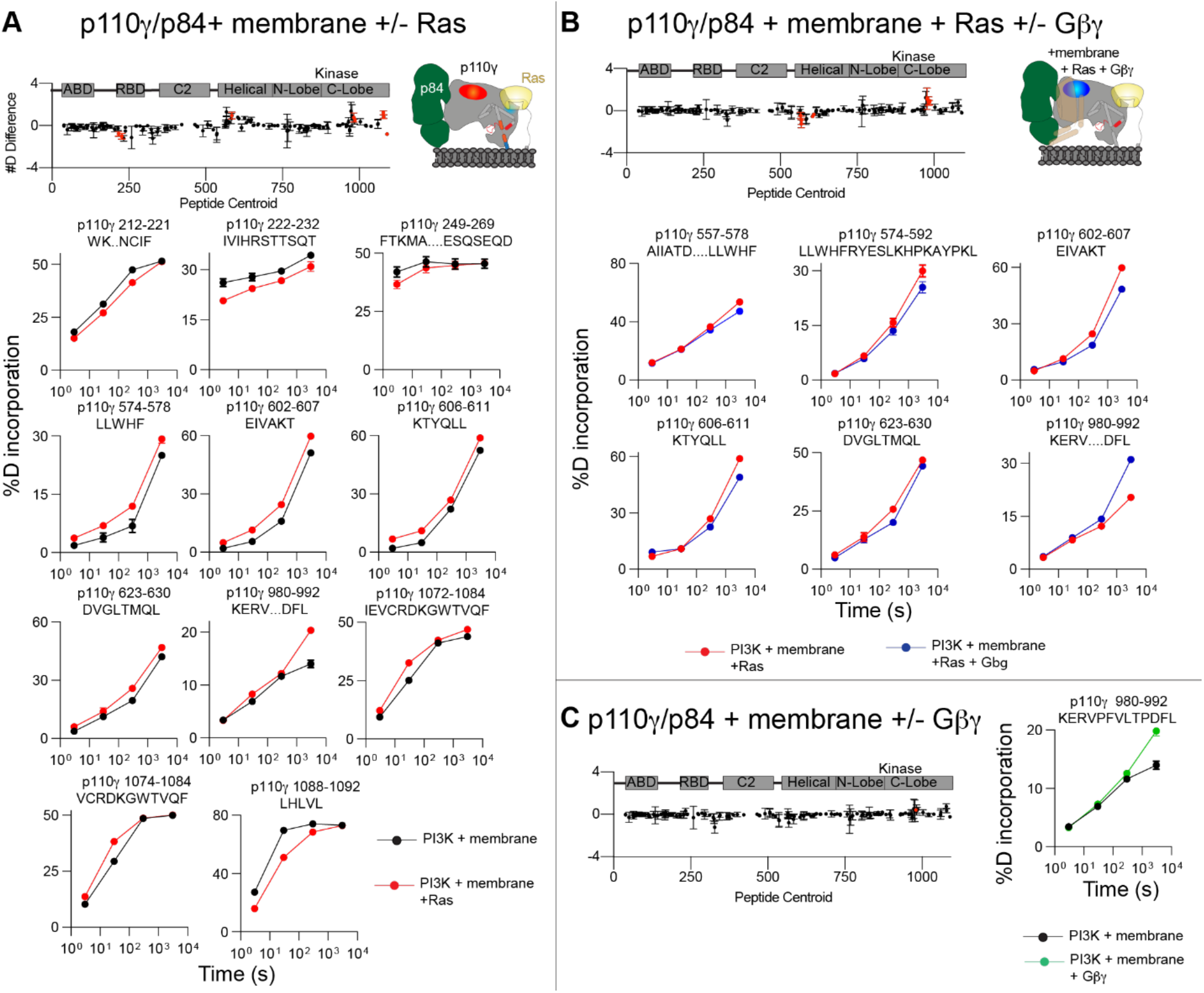
**(A-C).** The sum of the number of deuteron differences in the p110γ subunit between plasma membrane mimic vesicles and (A) plasma membrane mimic vesicles with 3 μM GTPγS loaded lipidated Hras and (B) plasma membrane mimic vesicles with 3 μM GTPγS loaded lipidated Hras 3 μM Gβγ and (C) plasma membrane mimic vesicles with 3 μM Gβγ. Each point is representative of the centre residue of an individual peptide. For all number of deuteron difference graph the peptides that met the significance criteria are coloured red, with error shown as standard deviation (n=3). A cartoon model is shown to the right with differences annotated. Selected deuterium exchange incorporation curves for peptides in the presence and absence of HRas and or Gβγ are shown below and are coloured according to the legend. Error is shown as standard deviation (n=3).

**Fig S5.**
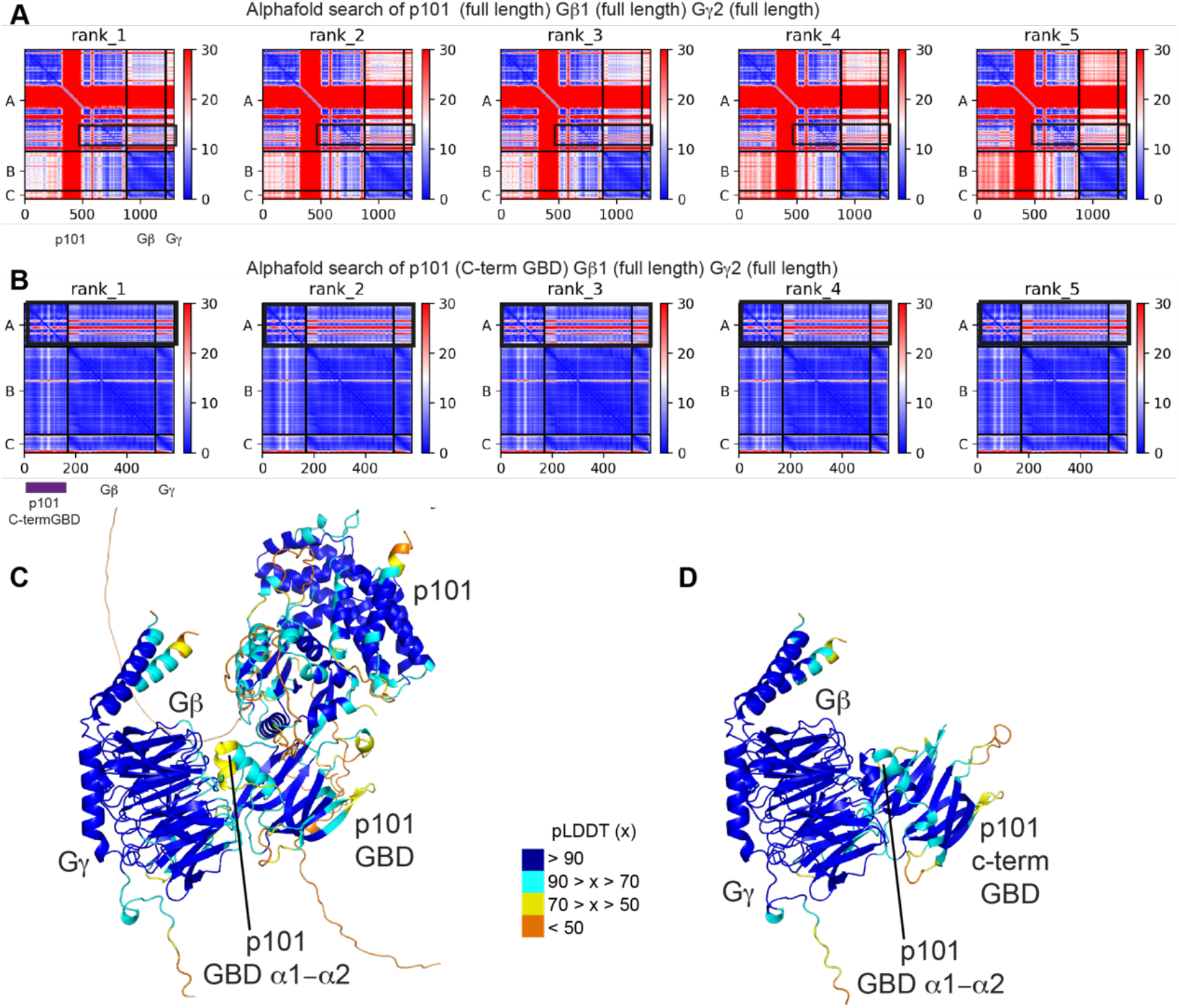
Alphafold2 multimer modelling of the complex between p101 and Gβγ. **A.** Predicted aligned error (PAE) for Alphafold2 Multimer search of full length p101, Gβ1, and Gγ2. The sequence of the two searches are indicated. **B.** Predicted aligned error (PAE) for Alphafold2 Multimer search of the C-terminus of p101, Gβ1, and Gγ2. The sequence of the two searches are indicated, with a schematic indicated below. For both panels A+B, the colours indicate the predicted aligned error, and are coloured according to the legend. Note that the PAE plot is not an inter-residue distance map or a contact map. Instead, the red-blue colour indicates expected distance error. The colour at (x, y) corresponds to the expected distance error in residue x’s position, when the prediction are aligned on residue y (more information can be found at https://alphafold.ebi.ac.uk/) (*37, 66*). Blue is indicative of low PAE, with the low PAE at the p101-Gβγ interface in both panels A+B suggests that AlphaFold2-multimer predicts the relative positions with high accuracy. **C+D.** Alphafold2 models from panels **A+B** shown with the per-residue confidence metric predicted local-distance difference test (pLDDT) coloured according to the legend. The pLDDT score varies from 0 to 100, and is an estimate of how well the prediction would agree with an experimental structure based on the local distance difference test Cα (*37*).

**Fig S6.**
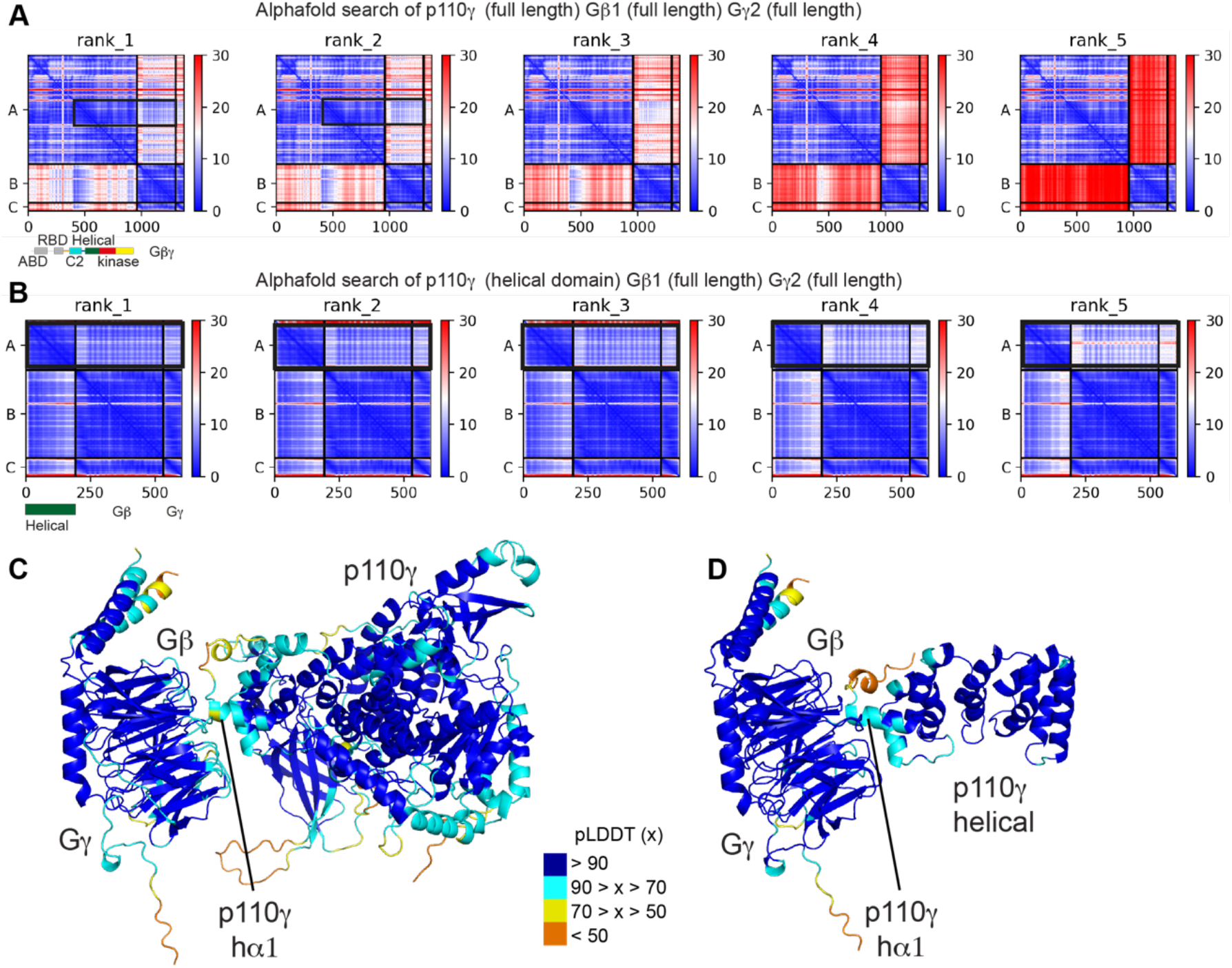
Alphafold2 multimer modelling of the complex between p110γ and Gβγ. **A.** Predicted aligned error (PAE) for Alphafold2 Multimer search of full length p110γ, Gβ1, and Gγ2. The sequence of the two searches are indicated, with a schematic indicated below. **B.** Predicted aligned error (PAE) for Alphafold2 Multimer search of the helical domain of p110γ, and full length Gβ1, and Gγ2. The sequence of the two searches are indicated, with a schematic indicated below. For both panels A+B, the colours indicate the predicted aligned error, and are coloured according to the legend. Note that the PAE plot is not an inter-residue distance map or a contact map. Instead, the red-blue colour indicates expected distance error. The colour at (x, y) corresponds to the expected distance error in residue x’s position, when the prediction are aligned on residue y (more information can be found at https://alphafold.ebi.ac.uk/) (*37, 66*). Blue is indicative of low PAE, with the low PAE at the helical domain-Gβγ interface in both panels A+B suggests that AlphaFold2-multimer predicts the relative positions with high accuracy. **C+D.** Alphafold2 models from panels **A+B** shown with the per-residue confidence metric predicted local-distance difference test (pLDDT) coloured according to the legend. The pLDDT score varies from 0 to 100, and is an estimate of how well the prediction would agree with an experimental structure based on the local distance difference test Cα (*37*).

**Table S1.** X-ray Data collection and refinement statistics.

|  | p110γ p84 |
| --- | --- |
| **Data collection** |  |
| Wavelength | 0.9762 |
| Space group | C121 |
| Cell dimensions |  |
| *a*, *b*, *c* (Å) | 322.3, 166.6, 255.8 |
| $\alpha$, $\beta$, $\gamma$ (°) | 90, 114, 90 |
| Resolution (Å) | 90.3 - 8.5 (8.89-8.5)* |
| *R*<sub>merge</sub> | 0.122 (0.39) |
| *I* / $\sigma I$ | 6.8 (4.4) |
| CC1/2 | 0.99 (0.90) |
| Completeness (%) | 99.9% |
| Redundancy | 5.8 |
| **Refinement** |  |
| Resolution (Å) | 90.3 - 8.5 (8.89-8.5) |
| No. unique reflections | 10833 (1084) |
| *R*<sub>work</sub> / *R*<sub>free</sub> | 27.9/33.9 |
| No. atoms |  |
| Protein | 50,008 |
| *B*-factors |  |
| Protein | 545.9 |
| Clash score | 5.72 |
| Ramachandran favored | 98.29 |
| Ramachandran outliers | 0.07 |
| Rotamer outliers | 0.50 |
| R.m.s. deviations |  |
| Bond lengths (Å) | 0.004 |
| Bond angles (°) | 0.64 |

\*Values in parentheses are for highest-resolution shell.
Number of crystals used for structure=1

**Table S2.**
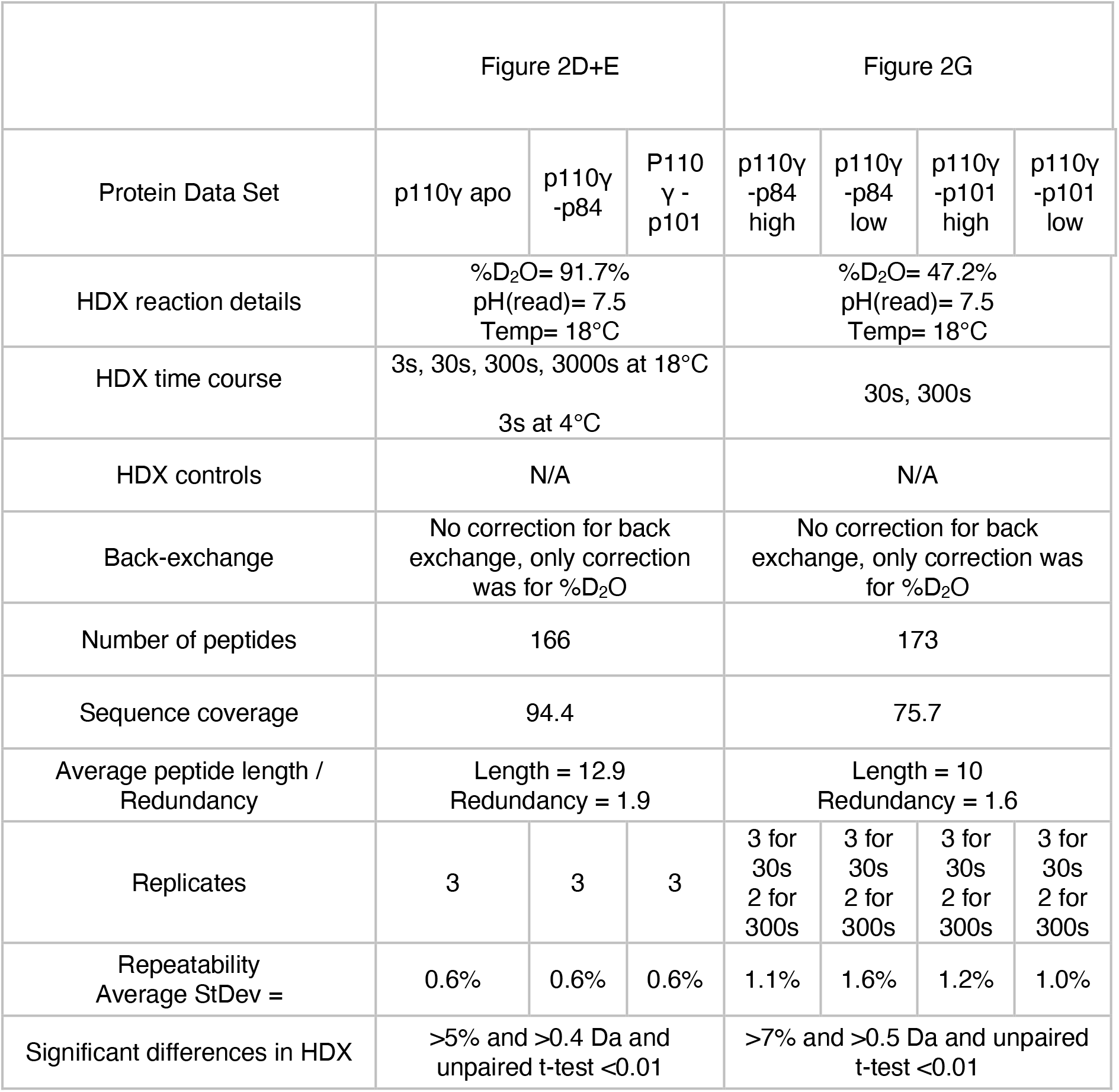
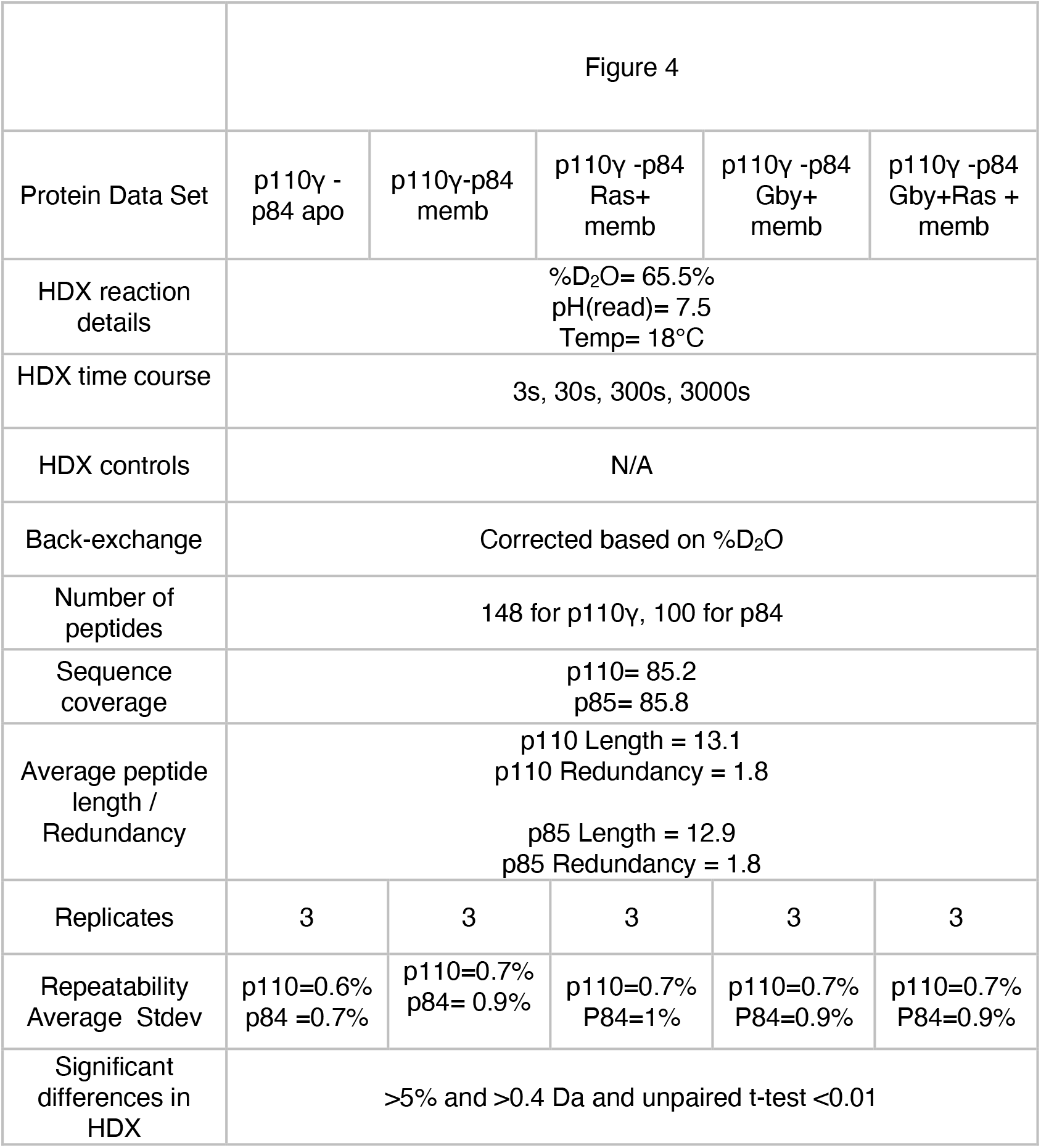
HDX-MS data processing table.

|  | Figure 2D+E |  |  | Figure 2G |  |  |  |
| --- | --- | --- | --- | --- | --- | --- | --- |
| Protein Data Set | p110γ apo | p110γ -p84 | P110 γ -p101 | p110γ -p84 high | p110γ -p84 low | p110γ -p101 high | p110γ -p101 low |
| HDX reaction details | %D<sub>2</sub>O= 91.7%<br>pH(read)= 7.5<br>Temp= 18°C |  |  | %D<sub>2</sub>O= 47.2%<br>pH(read)= 7.5<br>Temp= 18°C |  |  |  |
| HDX time course | 3s, 30s, 300s, 3000s at 18°C<br>3s at 4°C |  |  | 30s, 300s |  |  |  |
| HDX controls | N/A |  |  | N/A |  |  |  |
| Back-exchange | No correction for back exchange, only correction was for %D<sub>2</sub>O |  |  | No correction for back exchange, only correction was for %D<sub>2</sub>O |  |  |  |
| Number of peptides | 166 |  |  | 173 |  |  |  |
| Sequence coverage | 94.4 |  |  | 75.7 |  |  |  |
| Average peptide length / Redundancy | Length = 12.9<br>Redundancy = 1.9 |  |  | Length = 10<br>Redundancy = 1.6 |  |  |  |
| Replicates | 3 | 3 | 3 | 3 for 30s<br>2 for 300s | 3 for 30s<br>2 for 300s | 3 for 30s<br>2 for 300s | 3 for 30s<br>2 for 300s |
| Repeatability<br>Average StDev = | 0.6% | 0.6% | 0.6% | 1.1% | 1.6% | 1.2% | 1.0% |
| Significant differences in HDX | >5% and >0.4 Da and unpaired t-test <0.01 |  |  | >7% and >0.5 Da and unpaired t-test <0.01 |  |  |  |

**Table S2. HDX-MS data processing table (cont.)**
|  |  |  |  |  |  |
| --- | --- | --- | --- | --- | --- |
|  | Figure 4 |  |  |  |  |
| Protein Data Set | p110γ -<br>p84 apo | p110γ-p84<br>memb | p110γ -p84<br>Ras+<br>memb | p110γ -p84<br>Gby+<br>memb | p110γ -p84<br>Gby+Ras +<br>memb |
| HDX reaction<br>details | %D<sub>2</sub>O= 65.5%<br>pH(read)= 7.5<br>Temp= 18°C |  |  |  |  |
| HDX time course | 3s, 30s, 300s, 3000s |  |  |  |  |
| HDX controls | N/A |  |  |  |  |
| Back-exchange | Corrected based on %D<sub>2</sub>O |  |  |  |  |
| Number of<br>peptides | 148 for p110γ, 100 for p84 |  |  |  |  |
| Sequence<br>coverage | p110= 85.2<br>p85= 85.8 |  |  |  |  |
| Average peptide<br>length /<br>Redundancy | p110 Length = 13.1<br>p110 Redundancy = 1.8<br>p85 Length = 12.9<br>p85 Redundancy = 1.8 |  |  |  |  |
| Replicates | 3 | 3 | 3 | 3 | 3 |
| Repeatability<br>Average Stdev | p110=0.6%<br>p84 =0.7% | p110=0.7%<br>p84= 0.9% | p110=0.7%<br>P84=1% | p110=0.7%<br>P84=0.9% | p110=0.7%<br>P84=0.9% |
| Significant<br>differences in<br>HDX | >5% and >0.4 Da and unpaired t-test <0.01 |  |  |  |  |

